# Glial dysregulation in human brain in Fragile X-related disorders

**DOI:** 10.1101/2022.03.29.486195

**Authors:** Caroline M. Dias, Maya Talukdar, Shyam K. Akula, Katherine Walsh, Christopher A. Walsh

## Abstract

While large trinucleotide repeat expansions at the *FMR1* locus cause Fragile X Syndrome (FXS), smaller “premutations” are associated with the late-onset condition Fragile X-associated tremor/ataxia syndrome (FXTAS), which shows very different clinical and pathological features, with no clear molecular explanation for these marked differences. One prevailing theory posits that the premutation uniquely causes neurotoxic increases in FMR1 mRNA (i.e., 4-8-fold increases), but evidence to support this hypothesis is largely derived from analysis of peripheral blood. We applied single-nucleus RNA-sequencing to post-mortem frontal cortex and cerebellum from 9 individuals with Fragile X mutations as well as age and sex matched controls (n=6) to assess cell-type specific molecular neuropathology. We found robust reduction of FMR1 mRNA in FXS as expected, with modest but significant upregulation (∼1.3 fold) of FMR1 in glial clusters associated with premutation expansions. In premutation cases we identified alterations in glia number in cortex and cerebellum. Differential expression analysis demonstrated altered cortical oligodendrocyte development, while gene ontology analysis revealed alterations in neuroregulatory roles of glia, such as glial modulation of neurotransmission and synaptic structure. We identified significant enrichment of known FMR1 protein target genes in differentially expressed gene lists in FXS as well as the premutation, suggesting FMR1 protein target pathways may represent a shared source of dysfunction in both conditions despite opposite FMR1 mRNA changes. These findings challenge existing dogma regarding FXTAS and implicate glial dysregulation as a critical facet of premutation pathophysiology, representing novel therapeutic targets directly derived from the human condition.

## Introduction

*FMR1* related disorders contribute to neurologic dysfunction across the lifespan (Leehey, 2009; Hagerman et al., 2017). Large trinucleotide (CGG) expansion (i.e., full mutations, FM) in the 5’ UTR of the *FMR1* gene are associated with the neurodevelopmental disorder Fragile X Syndrome (FXS) while smaller, premutations (PM) are associated with Fragile X-associated tremor and ataxia syndrome (FXTAS), a late onset condition characterized by executive functioning decline and progressive cerebellar ataxia, presenting in 40-70% of PM carriers (Fu et al., 1991; Pieretti et al., 1991; Verkerk et al., 1991; Jacquemont et al., 2003; Grigsby et al., 2016). In the latter, neuropathological and imaging studies have identified intranuclear neuronal and astrocytic inclusions, prominent white matter abnormalities including myelin pallor and spongiosis, and characteristic T2 white matter hyperintensities on MRI (Jacquemont et al., 2003; Tassone et al., 2004; Cohen et al., 2006; Greco et al., 2006; Schwartz et al., 2021). In contrast, in Fragile X syndrome, an early-onset neurodevelopmental disorder characterized by intellectual disability, autistic symptoms and characteristic facial features (Martin & Bell, 1943; Turner et al., 1975), only subtle functional changes in white matter in humans have been identified on imaging (Hallahan et al., 2011; Sandoval et al., 2018; Swanson et al., 2018). The molecular correlates of these white matter findings in both conditions are unknown.

The FM is associated with hypermethylation and transcriptional silencing of the *FMR1* locus, and absence of FMR1 protein (FRMP), while the PM has been reported to be paradoxically associated with increases in FMR1 mRNA, particularly in blood, with only variable reductions in FMRP levels (Fu et al., 1991; Oberle et al., 1991; Pieretti et al., 1991; Verkerk et al., 1991; Tassone et al., 1999; Tassone, Hagerman, Chamberlain, et al., 2000; Tassone, Hagerman, Taylor, et al., 2000; Kenneson et al., 2001). Although individuals with the PM may present with alterations in typical neurodevelopment, FXS patients generally do not present with features of FXTAS. These divergent clinical and molecular phenotypes have led to the hypothesis that the clinical symptomatology associated with FXTAS is related to a neurotoxic effect of increased levels of FMR1 mRNA in the nervous system. This argument is bolstered by findings of a 4-8 fold increase of FMR1 mRNA in peripheral blood cells of individuals with the PM (Tassone, Hagerman, Taylor, et al., 2000). However, prior bulk studies of human post-mortem brain tissue from individuals with the PM have revealed more modest ∼0.9-1.5 fold changes in FMR1 mRNA (Tassone et al., 2004; Pretto et al., 2014). Prior studies in post-mortem human brain in both FXS and FXTAS have focused on bulk cellular analysis, which does not resolve cell-type specific molecular alterations. Whereas it is possible that the cellular heterogeneity of the human CNS may mask toxic levels of FMR1 mRNA, other hypotheses, including an inappropriate DNA damage response, mitochondrial stress, and polyglycine-containing peptide accumulation, have been put forth as alternative hypotheses to explain the pathophysiology of FXTAS (Garcia-Arocena & Hagerman, 2010; Sellier et al., 2017; Schwartz et al., 2021). It is also possible that reduced FMRP contributes to PM pathology in a developmentally distinct manner from the total loss that occurs in FXS. Finally, while studies of the impact of *FMR1* disruption have been focused on post-mitotic neurons, there is increasing evidence implicating important roles for *FMR1* in a diversity of cellular subtypes at multiple points in nervous system development, including in glia (Giampetruzzi et al., 2013; Martínez Cerdeño et al., 2018; Doll et al., 2020; Doll et al., 2021; Raj et al., 2021).

Despite these gaps in knowledge, there has been no cell-type specific analysis of transcriptional changes related to Fragile X in human brain to date.

To understand the molecular and cellular perturbations associated with Fragile X expansion in human brain across the lifespan with an unbiased approach, we applied single nuclei RNA-sequencing (snRNA-seq) to post-mortem frontal cortex and cerebellar hemisphere of individuals with *FMR1* PMs and FXS. We identified changes in *FMR1* expression, cellular proportion, and global gene expression that challenge current assumptions about molecular mechanisms underlying FXTAS pathogenesis, and specifically implicate glial dysregulation as critical in Fragile X molecular neuropathology.

## Results

Prior to tissue processing we reviewed available medical records to ensure that clinical and neuropathological data were consistent with genetic diagnoses (Table 1, Methods). All cases here have been previously presented in prior published work (Lohith et al., 2013; Esanov et al., 2016; Tran et al., 2019). Although we focus on the PM, we also included two known cases of FXS, to assess whether well-known effects on *FMR1* expression were present in our dataset. One case of FXS due to a deletion of *FMR1* was included given the known shared molecular consequences of *FMR*1 deletion and trinucleotide expansion (Gedeon et al., 1992). Neither FXS case had neuropathological abnormalities noted, consistent with expectations. The majority of PM cases had either clinical and/or neuropathological evidence of FXTAS. We identified one case that in the past was mistakenly categorized as FXS, but whose clinical records and genetic testing revealed it to be a PM (see Table 1). We validated functional effects of Fragile X disruption with western blotting of FMRP directly on frontal cortex tissue (Figure 1A) and obtained expected results: absent FMRP in FXS in both the FM and gene deletion, and variably reduced FMRP in PM cases.

**Figure 1:**
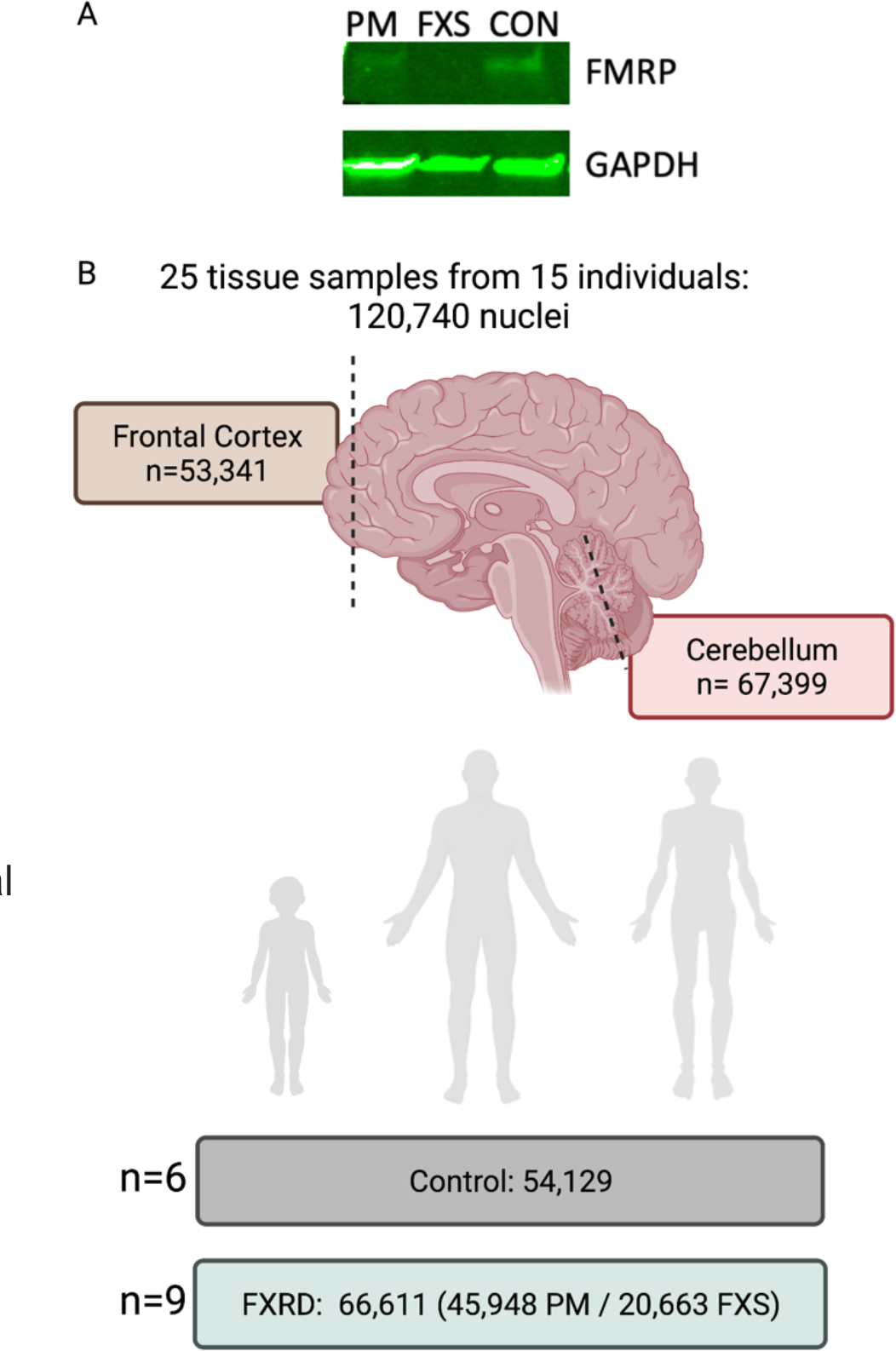
Tissue validation and sample size overview. A. Representative western blot of frontal cortex protein lysate demonstrating absent FMRP in FXS and variably reduced FMRP in PM case. B. Summary of sample size and final filtered nuclei number per condition and region, representing samples from across the lifespan in all conditions. FXRD: Fragile X related disorders.

**Table 1:**
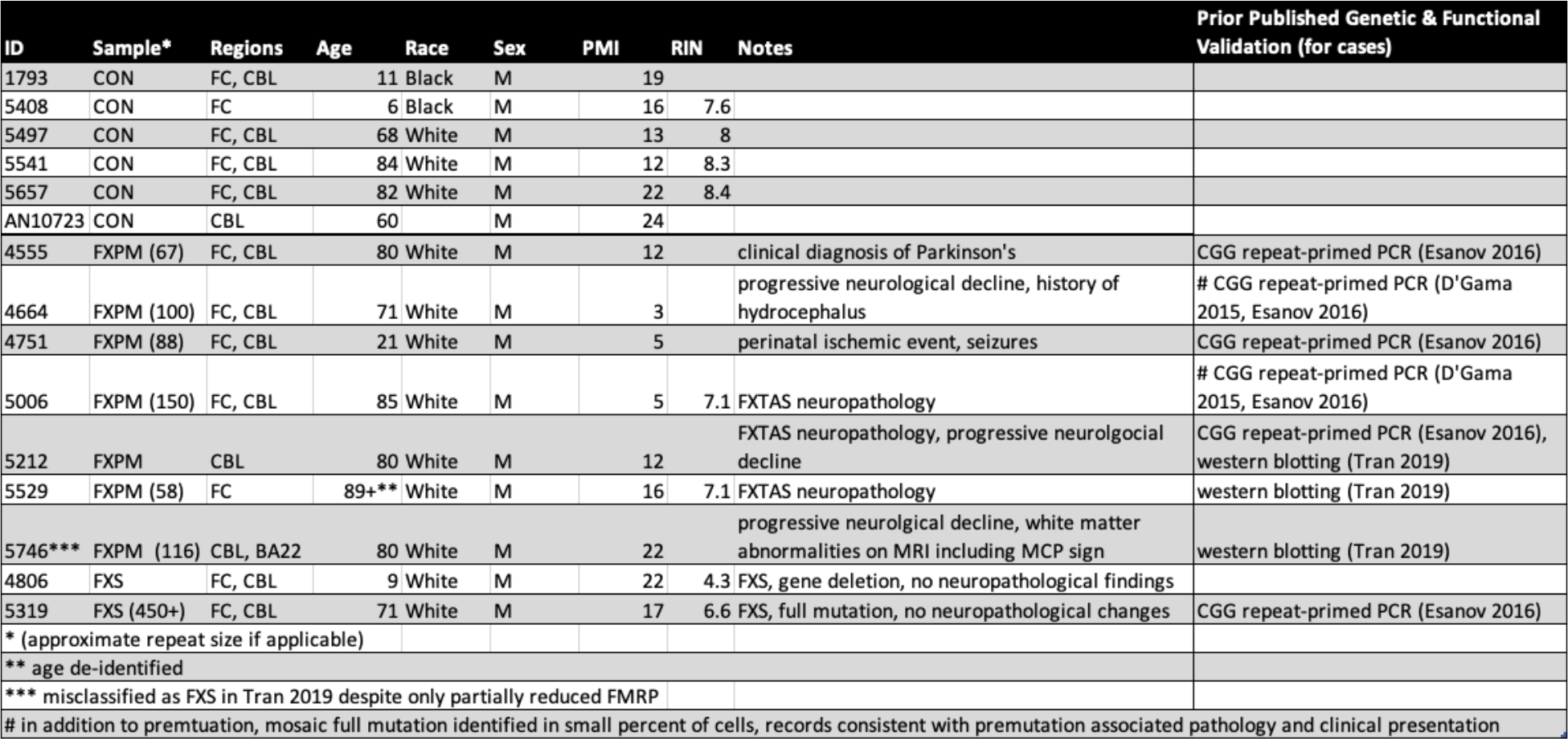
Demographic information for post-mortem samples used. Repeat size if applicable ascertained from clinical records and prior published work. FC: frontal cortex, CBL: cerebellum

Following unsupervised clustering of single nuclei, and filtering, we obtained over 120,000 high quality nuclei for further analysis across samples, including nuclei from 6 age- and sex-matched controls (Figure 1B, Table 2). We applied known cell type-specific markers to assess the specificity and accuracy of unsupervised clustering (Fig 2). For both prefrontal cortex and cerebellar hemisphere we identified specific classification of cellular subtypes for both neurons and glia. To further validate our annotation, we also identified layer-specific excitatory neurons and inhibitory neuron subclusters in the frontal cortex which were consistent with broader categorizations and prior annotations (Figure 2 figure supplement 1) (Lake et al., 2016; Hodge et al., 2019; Velmeshev et al., 2019; Langseth et al., 2021) We chose to group excitatory subtypes, and group inhibitory subtypes respectively, to maximize power for downstream analysis.

**Figure 2:**
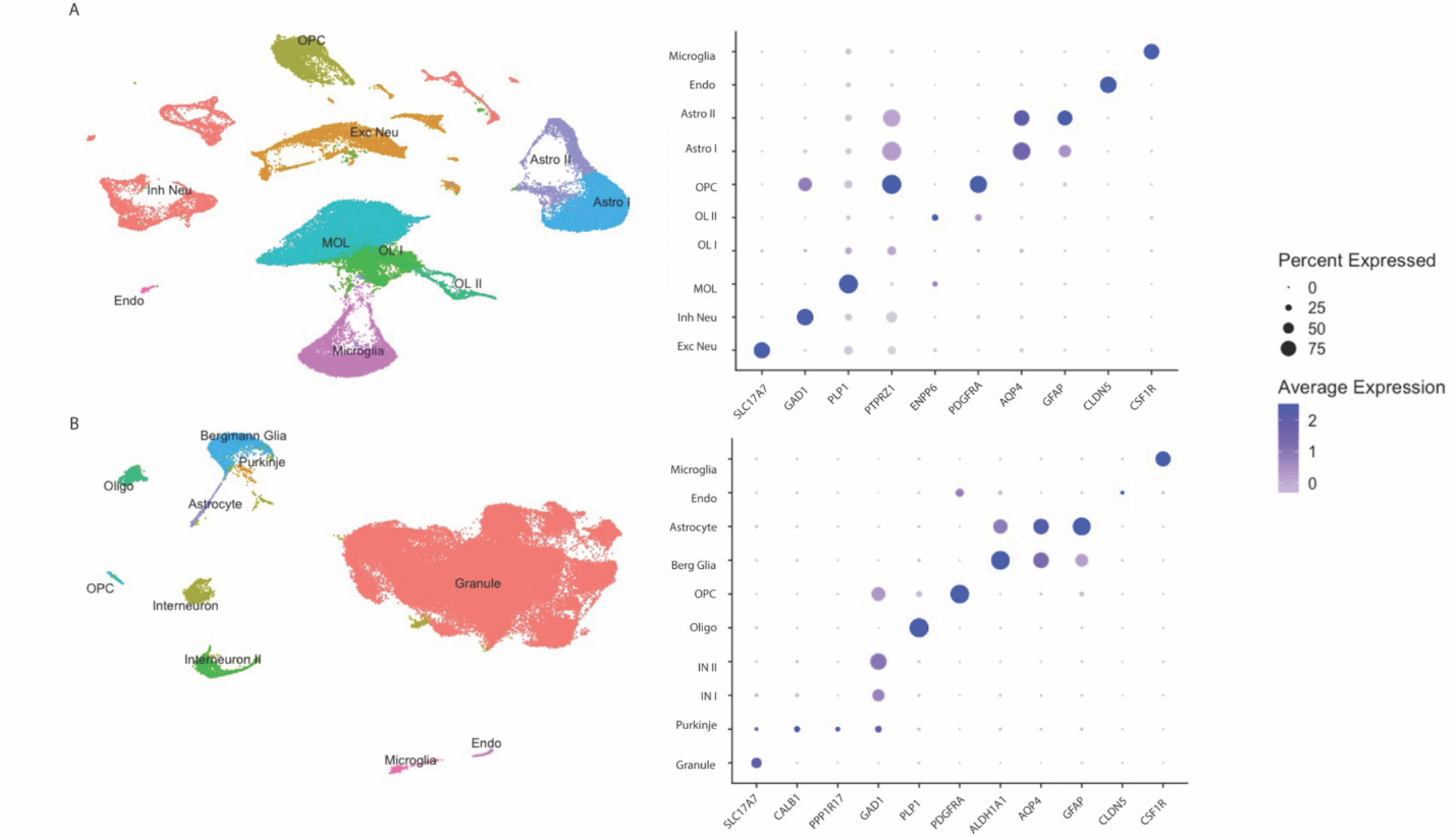
snRNA-seq of frontal cortex and cerebellum. Unsupervised clustering and visualization in Seurat visualized with UMAP (left) as well dot plot of cell type specific gene expression (right) in frontal cortex (A) and cerebellum (B) reflects accurate and specific cell type classification. Sample size as in Figure 1. Abbreviations: Astro: astrocyte, endo: endothelial, OPC: oligodendrocyte progenitor, OL: oligodendrocyte lineage, MOL: mature cortical oligodendrocyte, Inh: inhibitory, Exc: excitatory, Berg glia: Bergmann glia, Oligo: mature cerebellar oligodendrocyte, IN: cerebellar interneuron.

**Table 2:** Filtered nuclei number by brain region and cluster status.

| Cluster | CON | FXPM | FXS | Total |
| --- | --- | --- | --- | --- |
| **Cerebellum** |  |  |  |  |
| Granule | 25067 | 24237 | 9831 |  |
| Oligo | 366 | 542 | 130 |  |
| Bergmann Glia | 643 | 1276 | 334 |  |
| Interneuron | 497 | 848 | 155 |  |
| Interneuron II | 395 | 930 | 97 |  |
| Microglia | 219 | 268 | 85 |  |
| Astrocyte | 263 | 276 | 104 |  |
| OPC | 157 | 327 | 79 |  |
| Endothelial | 87 | 63 | 20 |  |
| Purkinje | 59 | 21 | 23 |  |
| **Cerebellar Total** | 27753 | 28788 | 10858 | 67399 |
| **Cortex** |  |  |  |  |
| Inh Neu | 3916 | 2049 | 1081 |  |
| Exc Neu | 2776 | 2689 | 868 |  |
| OPC | 2411 | 1467 | 1591 |  |
| OL I | 3173 | 1058 | 522 |  |
| OL II | 250 | 81 | 630 |  |
| MOL | 4381 | 6890 | 1767 |  |
| Astro I | 4264 | 914 | 1577 |  |
| Astro II | 1719 | 468 | 614 |  |
| Microglia | 3348 | 1518 | 1092 |  |
| Endo | 138 | 26 | 63 |  |
| **Cortex Total** | 26376 | 17160 | 9805 | 53341 |

There were distinctions between cerebellum and cortex in overall glial cell composition. In frontal cortex we identified several distinct clusters appearing to reflect different states of oligodendrocyte development, including PDGFRA + oligodendrocyte progenitor cells (OPCs), two intermediate clusters (OLI-ENPP6+ and OLII-TFC7L2+), and a mature myelinating oligodendrocyte (MOL) cluster. We compared the transcriptional profile of OLI and OLII to oligodendrocyte lineage clusters identified in mouse (Marques et al., 2016), and found that OLI gene expression resembled mouse committed oligodendrocyte progenitors (COPs) and OLII resembled immature, newly formed, non-myelinating oligodendrocytes. On the other hand, in the cerebellum, although granule cells accounted for the majority of nuclei captured, as expected, we also identified a cerebellar specific Bergmann glia cluster. OLI and OLII clusters were not identified in cerebellar samples.

Comparison of average FMR1 mRNA across all nuclei in frontal cortex and cerebellum revealed regional differences in relative expression between neurons and glia. In the frontal cortex, expression of *FMR1* was higher in excitatory and inhibitory neurons compared to glia, as expected (Figure 2 figure supplement 2, Figure 3).

**Figure 3:**
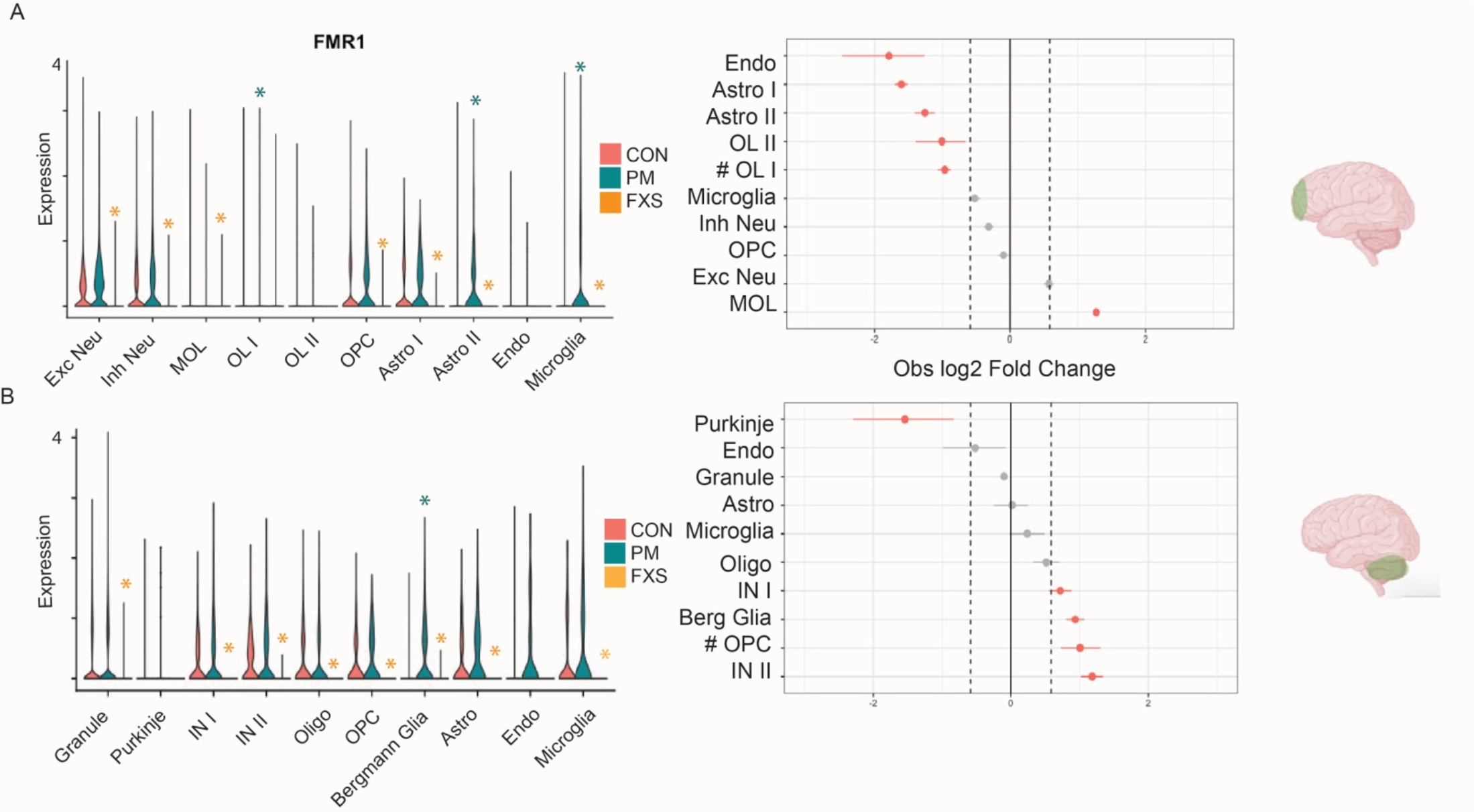
FMR1 mRNA dysregulation in FXS and PM cases, and cluster proportion analysis for PM cases. Frontal cortex (**A**) and cerebellar (**B**) changes in FMR1 expression in FXS and PM (left) and cellular proportion alterations in PM cases vs control (right) A. Violin plots demonstrate reduced FMR1 mRNA expression in multiple clusters in FXS, while FMR1 mRNA is variably increased in PM clusters, primarily in glia. Cluster abbreviations as in Fig. 2. Orange *: reduced FMR1 p-value in FXS vs CON padj< .05, teal *: increased FMR1 in PM vs. CON padj < .05. Right panel shows permutation plot for PM cluster proportions demonstrating glial cell number alterations in cortex and cerebellum. Red: FDR < .05 and abs(log2FC) > 0.58. # indicates significance of this cluster is not robust to outlier sample removal.

However, in the cerebellum, *FMR1* expression was expressed in more non-neuronal subtypes at baseline. We used an independent harmonized single cell transcriptomic resource to confirm these findings and identified similar region-specific patterns of expression with higher relative glial to neuron FMR1 mRNA expression in cerebellum. (Figure 2 figure supplement 3) (Song et al., 2021).

Analysis of individual cluster *FMR1* expression revealed cell type-specific effects of Fragile X status on *FMR1* transcription. In both cases of Fragile X syndrome, we identified equal and total abrogation of *FMR1* expression, as expected (Figure 2 figure supplement 4). Although our sample of FXS cases is small, this provides important proof of principle that expected transcriptional signatures are present within the snRNA- seq data. Indeed, despite the smaller sample size and nuclei number, reduced *FMR1* expression was robust among different clusters in FXS in both neuronal and glial subpopulations across the brain, consistent with the large effect size of this genetic driver (Figure 2 figure supplement 4, Figure 3). The effect of the PM on FMR1 expression, despite the larger sample size, was more modest and demonstrated cluster-specific heterogeneity. In PM cases, the only clusters that demonstrated significantly increased FMR1 mRNA expression in either the frontal cortex or cerebellum were non-neuronal, such as cerebellar Bergmann glia and cortical microglia (Figure 3). Although the absence of significant FMR1 upregulation in a small number of clusters in PM cases may be due to inadequate power, most clusters included more than enough nuclei (including frontal cortex excitatory and inhibitory neurons, as well as cerebellar granule cells and interneurons) to rule this explanation out (see Methods).

Thus, in general, the lack of significant upregulation of *FMR1* mRNA in neuronal subclusters in the PM cases is not due to a lack of power. Rather, it suggests that overall, the increase in FMR1 expression in brain caused by the PM is far more modest than the 4-8 fold increase observed in blood, and shows a preferential impact on glia, in the regions assessed here.

In the cerebellum, we identified changes in nuclei number in PM cases that recapitulated past neuropathological studies, specifically previous work demonstrating cerebellar Purkinje cell loss and Bergmann cell gliosis in individuals with the PM (Hagerman, 2013). Consistent with this, we identified fewer Purkinje cell nuclei, and greater Bergmann cell nuclei, in the cerebellum of PM carriers versus age matched controls (Figure 3). This orthogonal validation of prior findings reinforces the utility of our molecular approach to identify bonafide biological phenomenon.

We also identified changes in glial cell number in the frontal cortex (primarily BA10) in association with the PM. PM cases demonstrated fewer than expected astrocytes and endothelial cells, with an increase in mature oligodendrocytes compared to controls, observations surprisingly consistent across samples and not reflecting the presence of outliers (Figure 3, Methods). We assessed BA22 in one PM sample and identified MOL levels and astrocyte levels to be similar to control, suggesting these findings may reflect a sub-cortical specific effect. Because changes in glial number have been observed in normal aging (Peters et al., 1991; Peters & Sethares, 2004; Soreq et al., 2017; Salas et al., 2020; Xu et al., 2020) we assessed the impact of age on these clusters and did not identify a significant association with age for these clusters in control individuals (Figure 3 figure supplement 1). We wondered whether age associated changes in neuronal composition however might mask more subtle effects of the PM on neuronal clusters, given that age-associated changes in interneuron density have also been observed (Hua et al., 2008; Stanley et al., 2012; Rozycka & Liguz-Lecznar, 2017) In this case, we identified a significant age-related decline in inhibitory neuron/total neuron composition in frontal cortex as previously reported (Majdi et al., 2007), although there was no detectable effect of Fragile X PM status on this decline (Figure 3 figure supplement 2). Thus, we identified unexpected novel alterations in glial number in frontal cortex in PM cases.

Although limited by small sample size, we also assessed cluster proportions in FXS and found that alterations in cellular composition was markedly distinct from PM cases. For example, frontal cortex astrocyte and endothelial cell depletion were not identified in FXS. (Figure 3 figure supplement 3). In cerebral cortex in FXS there was an increase in OPC proportion; this finding has previously been observed in animal models (Doll et al., 2021) although to our knowledge not previously described in humans. Thus, FXS and PM cases are associated with distinct alterations in the brain’s cellular architecture.

In frontal cortex, we identified alterations in glial cell transcriptional regulation indicating widespread perturbations in PM pathology (Figure 4). We focused on the frontal cortex given the broader and more equitable distribution of cell types, which allows for more direct comparisons between clusters. In PM cases, we identified a marked trend towards increased expression of myelination markers and reduced immature markers in the intermediate committed oligodendrocyte progenitor OLI cluster (Figure 4A). The increase in mature markers in an intermediate progenitor offers a possible explanation for the increased number of MOLs in PM frontal cortex despite no change in the OPC pool number. Indeed, committed oligodendrocyte progenitors are known to serve as a reservoir for rapidly generating myelinating mature oligodendrocytes (Lee et al., 2012; Lecca et al., 2020). The OLI nuclei cluster was not the only cluster with alterations in critical glial developmental gene expression in PM cases. For example, within PM OPCs, upregulation of CSPG4 (i.e., NG2) (Ampofo et al., 2017) was observed in frontal cortex (Figure 4 supplemental figure 1). Intriguingly, FXS clusters also demonstrated dysregulation of several of glial markers, but with distinct directionality and magnitude of changes, and preferential impact on OPCs.

**Figure 4:**
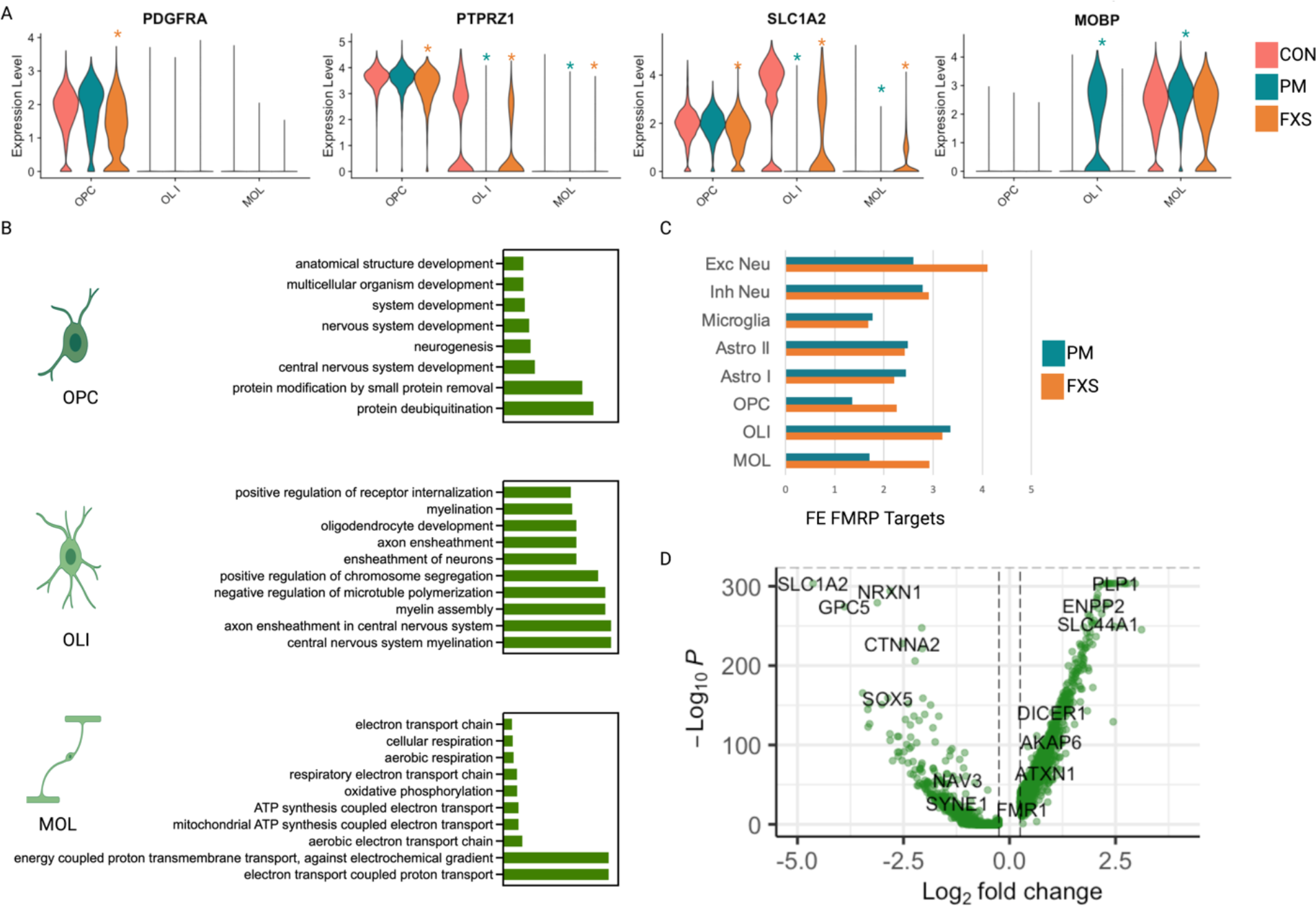
Global glial transcriptional dysregulation in frontal cortex. A. Violin plots demonstrating alteration in oligodendrocyte development in PM and FXS. Orange * indicates FXS vs. CON padj < .05, teal * indicates PM vs CON padj < .05. B. GO analysis of upregulated differentially expressed genes in oligodendrocyte clusters in frontal cortex demonstrate increased metabolic stress in MOLs and upregulation of myelination in OLI. C. Significant fold enrichment (FE) of FMRP target genes across clusters and conditions, all comparisons padj <.05 except for OPC in PM cases which was non-significant. D. Volcano plot of differentially regulated genes in PM cases in OLI demonstrate 3:1 increased: decreased DEGs, genes of potential interest noted.

There are thus cortical/cerebellar and condition specific patterns to glial transcriptional dysregulation in Fragile X related disorders.

Gene ontology analysis (see Methods, Figure 4) suggested that individual gene expression changes represented broader disturbances in biological functioning.

Interestingly, in PM cases, frontal cortex MOLs demonstrated unique evidence of increased metabolic stress, consistent with neuropathological findings of oligodendrocyte dysfunction. The OLI cluster in PM cases was the only group in either condition or region to demonstrate myelination upregulation (Figure 4B), demonstrating that the gene expression changes described above were specific to this cell cluster.

This is of particular interest given that several myelination related genes are known FMRP targets (Wang et al., 2004; Darnell et al., 2011; Giampetruzzi et al., 2013) - yet, these were not ubiquitously impacted in all oligodendrocyte lineage clusters.

Downregulated categories in PM cases implicated dysfunction of glial regulatory roles. For example, there was evidence of downregulation of synaptic organization and activity, decreased neuronal cell adhesion, and downregulation of glutamatergic signaling in a variety of oligodendrocyte lineage clusters (Figure 4 figure supplement 2). Inspection of biological processes disturbed in PM cases in astrocytes and microglia similarly revealed downregulation of several neuronal regulatory roles as well as evidence of inflammation (Figure 4 figure supplement 3). There was also evidence of neuronal dysfunction: neurons in PM cases demonstrated evidence of metabolic stress and downregulation of critical neuronal function and structure, such as postsynaptic organization and dendritic spine morphogenesis (Figure 4 figure supplement 4).

Inspection of differentially regulated genes in FXS revealed widespread evidence of increased metabolic and energetic stress, across both neurons and glia with known FMRP targets present among differentially regulated genes (Fig 4 figure supplements 5-8).

Given that we observed the presence of FMRP targets in both neuronal and glial differentially expressed gene lists in both FXS and PM cases, we investigated whether this reflected statistically significant enrichment (Figure 4C). We had hypothesized that there would be significant enrichment of FMRP target genes in FXS but not PM cases, and greater enrichment in neuronal clusters vs non-neuronal clusters. To our surprise, we identified significant enrichment for FMRP targets among differentially expressed genes across many cell clusters in both FXS and PM cases. Excitatory neurons demonstrated the highest fold enrichment of known FMRP targets in FXS as expected, but significant enrichment was observed in non-neuronal clusters in FXS as well.

Additionally, there was significant enrichment of FMRP target genes in differentially expressed genes across most clusters in the PM cases, in many cases very similar to FXS, suggesting a role for FMRP dysfunction in PM pathogenesis. The highest enrichment for FMRP targets among PM clusters was not neuronal but rather the OL1 cluster. Interestingly, this cluster exhibited global transcriptional dysregulation similar to FMRP loss in FXS in excitatory neurons including ∼ 3:1 increased:decreased significant differentially expressed genes (Fig 4D, Figure 4, figure supplement 9), reflecting a de- repressed state. However, known FMRP targets were not restricted to the upregulated list in this cluster (Figure 4D), potentially highlighting dual roles of FMRP as transcriptional repressor and in mRNA stabilization (Hale et al., 2021).

## Discussion

We present the first cell type specific analysis of gene expression of Fragile X related disorders in human brain. We identified changes in FMR1 mRNA expression, cellular proportion, and cell-type specific gene expression that sheds light on molecular perturbations associated with *FMR1* and specifically highlights an important role for glial molecular dysregulation in PM pathology.

Our data suggest that FMR1 mRNA expression in PM cases, at least in the brain regions analyzed here, is more modestly affected than has been observed in peripheral blood cells and furthermore that it preferentially effects glial cells more than neurons.

Given the robust elimination of FMR1 mRNA expression observed here in association with FXS, regardless of the genetic driver (trinucleotide expansion vs gene deletion), nuclei cluster, or brain region, we are confident in the validity of our approach. In the PM cases, we identified modest upregulation of *FMR1* expression, with glial subclusters in both cerebellum and cortex demonstrating the most marked increases. Rather than extreme neurotoxic increases in FMR1 mRNA, our findings suggest a more modest, ∼1.3 fold, increase in *FMR1* transcript levels, paralleling past studies of brain homogenate (Tassone et al., 2004; Pretto et al., 2014). Although we cannot rule out that neural cells expressing toxic levels of FMR1 transcript are selectively vulnerable and preferentially lost with time, our cellular proportion analysis (see below) does not support this interpretation, as one would expect clusters that are disproportionately lost to have relatively higher increases in *FMR1* mRNA. We also include in our analyses one 21 year-old PM case, whose data is very similar to the other aged PM cases, which further argues against age-related loss. Finally, changes in FMR1 expression were comparable between clusters known to be vulnerable to PM associated intranuclear inclusions (neurons, astrocytes) and those known to be spared (oligodendrocytes), arguing against inclusion presence as being a confounding factor in FMR1 mRNA measurement. Although our work challenges the causal role of extremely elevated FMR1 mRNA in human brain, it is possible that more modest increases of CGG containing FMR1 RNA still lead to cellular dysfunction, through previously posited mechanisms including trinucleotide repeat toxicity. Thus, our work provides an important foundation to understanding FMR1 mRNA levels that are relevant to neurological pathophysiology in animal and human model systems and broadens the scope of cellular subtypes that warrant further investigation within this context.

### Cellular proportion

Alterations in cell number in PM cases in both cerebellum and cortex also implicated glial dysregulation. Our finding of reduced Purkinje cells and Bergmann cell increases in the cerebellum in PM cases parallels well-described neuropathological findings (Greco et al., 2006), and reinforces the validity of our approach, and suggests that loss of Purkinje cells may contribute to FXTAS signs and symptoms. In PM cases, we also identified a proportional decrease in endothelial cells and astrocytes, with increases in MOLs in frontal cortex, a finding not explained by age or the presence of outliers. The relationship of basal FMR1 mRNA expression, change in FMR1 expression, and cellular proportion was not straightforward, arguing against a simplistic relationship between cellular proportion and FMR1 toxicity. For example, glial cells in the frontal cortex that demonstrated modest differential expression of FMR1 also demonstrated the most marked changes in cellular proportion. This may be related to earlier developmental time points that are impacted, cellular extrinsic effects on survival and proliferation, or both. Given the findings of global brain atrophy, white matter abnormalities, and the reported decline in executive functioning deficits reported in FXTAS (Brunberg et al., 2002; Greco et al., 2006; Brega et al., 2008; Grigsby et al., 2008), these changes in cellular proportion warrant further exploration of the role of glia in FXTAS associated cognitive decline and in other cortical areas.

The findings of changes in endothelial cell proportion are also interesting considering the recent description of microangiopathy in FXTAS neuropathology (Salcedo-Arellano et al., 2021). In fact, MRI findings in FXTAS of T2 hyperintensity have some similarities to microvascular ischemia (Leehey, 2009), also suggesting cerebrovascular dysfunction as an important facet of FXTAS. Past work has demonstrated that disruption of the blood brain barrier can lead to increased OPC NG2 expression (Rhodes et al., 2006), observed in our data in the frontal cortex in PM cases only. These findings highlight the need for additional work to interrogate the developmental mechanisms and regional specificity of *FMR1* effects on the endothelium on a more comprehensive level.

We identified global transcriptional alterations associated with PM status that support the now well-established principle that glia play central roles in neurodevelopment and disease (Teismann et al., 2003; Croisier & Graeber, 2006; Lee et al., 2012; Liu et al., 2015). For example, OPCs are known to form synaptic like structures and respond to neuronal activity (Bergles & Richardson, 2015) and glia more generally are critical in neuronal development, axonal integrity and behavior (Nave, 2010; Fernandes et al., 2017; Nagai et al., 2021). Differential gene regulation in PM glia frequently identified perturbations in glial-neuronal signaling, maintenance of synaptic structure and function, and altered neurotransmission in multiple glial lineage clusters in addition to evidence of energetic stress specifically in MOLs. Indeed, white matter abnormalities in Fragile X related disorders more broadly, may reflect subtle disruption of glial regulatory roles in neuronal homeostasis. Determination of whether these glial abnormalities contribute causally to clinical symptomatology or represent a secondary response to neuronal dysfunction, will require further work in human model systems.

Support for the former is corroborated by the fact that we identified significant enrichment of FRMP target genes in differentially expressed genes in both FXS and PM glial clusters. Although the magnitude of FMRP target enrichment was largest in excitatory neurons in FXS, there were surprising similarities in the magnitude of enrichment between FXS and PM. Although our work can not address causality, this suggests that shared loss of FMRP contributes to some degree to both disorders in a variety of cell types. There was also evidence of cell-type specific transcriptional alterations. For example, the OLI cluster, uniquely demonstrated significant changes in expression in myelin related genes (known FMRP targets) in PM cases, suggesting that cellular states even within a single glial lineage may be differentially vulnerable to Fragile X disruption in a context dependent manner. Thus, our findings in PM post-mortem brain, in light of known neuropathological and imaging abnormalities, support the interpretation of FXTAS as a disorder defined by glial dysfunction.

Our small sample size of Fragile X syndrome increases chances that changes in cellular proportion or transcriptional dysregulation may be due to stochastic artifacts. However, replication of past findings including reduced FMR1 expression (Pieretti et al., 1991; Bhattacharyya & Zhao, 2016), increased OPC proportion (Doll et al., 2021), and widespread evidence of metabolic stress (Donnard et al., 2020; Kang et al., 2021), corroborate known molecular neuropathology of the disorder (Licznerski et al., 2020).

Thus, our findings highlight the need for more comprehensive study of Fragile X in human tissue directly in a variety of different cell types.

In conclusion, we provide compelling evidence from human brain regarding cell type-specific molecular neuropathology that helps contextualize the clinical heterogeneity associated with genetic variation at the *FMR1* locus in neurodevelopment and neurodegeneration and specifically implicates glial dysregulation in PM pathology.

## Materials and Methods

### Samples

Post-mortem tissue was obtained from either the University of Maryland Neurobiobank and for one control case, from Autism Brain Net. All tissue was from deceased individuals and as such is not considered human subjects research. Fragile X mutation status/repeat size was determined through direct review of de-identified clinical records, and cross-referenced with prior published validation of the same cases (Table 1). Most of the PM cases had clinical symptomatology or neuropathological evidence of FXTAS (Table 1). Samples were matched for age, sex, and PMI but no cut-offs were utilized to exclude any cases (see Appendix Figure 1 for further details.)

### Western Blotting

∼25-50 mg frozen frontal cortex was homogenized in RIPA buffer + protease inhibitors and centrifuged, total protein content was then quantified. Laemelli sample buffer was added to the protein supernantant and boiled for 5 minutes. Equal amounts of protein (10 ug) were loaded onto precast SDS-Page gels with molecular weight ladders. Samples were transferred to membranes, blocked with Licor block (Lincoln, NE), and incubated in primary antibody overnight diluted in block at four degrees.

Following four washes in tris buffered saline + tween (TBS+T), blots were incubated with LI-COR secondary fluorescent antibodies in the dark at room temperature for one hour. After further washing including a final wash of TBS, the blots were scanned on a LI-COR Odyssey imager and images analyzed with Image-J software. The following primary antibodies and dilutions were used: GAPDH (Cell Signaling # 2118S) 1:15,000; FMRP (Cell Signaling 4317S) 1:1,000.

### Isolation of post-mortem nuclei

Frozen tissue (∼ 25 mg) from either section 1 frontal cortex or section 5 cerebellar hemisphere (see table of demographics) was dissected at -20 and subjected to dounce homogenization followed by sucrose gradient centrifugation as previously described. Nuclei were filtered and incubated for 5 minutes in 1:1000 Hoechst (Invitrogen H3569, Waltham MA). 10,000 Hoechst + nuclei from the suspension were then sorted directly into 10x Genomics RT buffer (Pleasanton, CA) on a chilled plate holder to remove doublets, debris, and dying nuclei on a FACS Aria (BD Biosciences, Franklin Lakes, NJ) with low pressure nozzle (Appendix Figure 2). Following sorting, reverse transcriptase enzyme was added on ice, and nuclei were immediately processed for encapsulation in the 10x Chromium controller. cDNA and libraries were prepared according to the 10x documentation protocol for 3’ gene expression. Libraries were prepared in matched batches and represent over 16 individual days of sample preparation and 4 separate sequencing runs.

### Sequencing and Quality Control

Following pooling of samples, pools were run on a Novaseq 6000 (Illumina, San Diego, CA) to obtain high coverage and saturation, and demultiplexed with bcl2fastq. CellRanger Count was utilized to generate count matrices with introns included, given intronic information is known to be informative for nuclear preparations. To remove remaining ambient RNA and debris, and obtain a final high quality nuclei set, filtering metrics were applied to nuclei in Seurat including: # UMIs> 500, # Genes > 250, log10GenesPerUMI (complexity measure) > 0.8, and mitoRatio < 0.1. Datasets were processed with SCTransform and integrated, and potential sources of variation were assessed with principal component analysis. Unsupervised clustering was performed with different resolutions followed by application of known cell type markers. For cerebellar Purkinjee and endothelial cells, which represented a small percentage of the total nuclei sample, cell type markers (CALB1; CLDN5) were used to manually select cell clusters using the SelectCells feature in Seurat.

### Analysis

Prism and R studio were used for analysis of demographic and single nuclei data, respectively, using statistical tests as indicated in each figure. Computationally intensive work was conducted on the Harvard Computing Cluster, O2. For analysis of demographic data, one PM case that was extremely aged is listed as 89+ to ensure sample de-identification and sample points were removed from any graphs presented here to ensure de-identification. For cluster proportion analysis, scProportion package (Miller et al., 2021) was used to perform a permutation test to calculate a p-value for each cluster, with a confidence interval returned via bootstrapping. (https://github.com/rpolicastro/scProportionTest). Although this approach is particularly amenable to small sample sizes, the outcome could still be impacted by samples with outlier cluster proportions. To assess the robustness of the data to the presence of outliers, cluster proportions from each individual sample were compared, and Grubb’s test used to identify the presence of outliers that may have contributed to significant results in scProportion. Samples which contributed to the presence of outliers were then systemically removed from the dataset and scProportion rerun. We identified only rare cases of outlier clusters that impacted significance of findings: One young control sample had a high proportion of the committed progenitor OL I. One PM sample demonstrated higher than expected OPC number. Other outliers did not impact significance of findings in PM cases and include: the FXS case with the gene deletion demonstrated an unusually high presence of the newly-formed, non-myelinating immature OLII cluster, and a different control sample had a higher proportion of endothelial cells. The plots presented represent the entire dataset with no data removed. Instead, cluster changes that were not robust to the removal of outlier samples are indicated on the graph with a # sign.

Differential expression analysis was done with the FindMarkers functionality in Seurat using default settings of Wilcoxon rank sum test, FDR < .05, log fold change > 0.25 for all cell clusters. A priori we calculated that 400 cells/condition cluster are required to detect 80% of differentially expressed genes with a false discovery rate of 5%. Thus, our analyses are adequately powered for the majority of cell cluster comparisons. We chose to omit downsampling to preserve power. Results were compared to a subset downsampled dataset for select clusters with larger nuclei number, and results were found to be similar both in the pattern of differentially regulated genes as well as the specific genes present in the data set (data not shown). Additionally, no association between cluster cell number and number of differentially expressed genes were identified. To generate a set of putative FMRP target genes in humans, the list from (Darnell et al., 2011) was transformed from mouse to human gene symbols resulting in a gene list of 745 predicted targets that were functionally validated in animal models. A hypergeometric test was performed for each cluster and condition, comparing the observed # FMRP target genes in differentially expressed lists vs predicted using a total gene set defined as 20,000. Gene ontology analysis was conducted with the web interface of Panther (http://geneontology.org) with default settings for statistical significance testing, including Fisher’s exact test and FDR < .05.

### Data and materials availability

All sequencing data and code are available from dbGaP (accession number pending).

## Appendix (Supplemental Methods)

We found no association between: PMI & RIN, age and PMI, and age and RIN, as expected (Appendix Figure 1). There was no difference in average RIN, PMI, or age between PM and controls. We did observe a reduction in RIN in the FXS samples as compared to controls but not PM cases.

Nuclear staining and sorting was conducted to select against dying cells, debris, and doublets (Appendix Figure 2).

## Acknowledgements

We thank Jennifer Neil for assistance with patient sample information; Robert Sean Hill, Dilenny Gonzalez and Sattar Khoshkoo for assistance in reagent ordering and sample sequencing; Sara Bizzoto and Sattar Khoskkoo for discussion of cell type specific markers in cortex, Ronald Mathieu and the Hematology/Oncology Flow Cytometry Research Facility for assistance with cell sorting; the Engle lab and the Harvard Biopolymers Facility for assistance with Chromium Controller use and sequencing. Molecular genetics library quantification support was provided by the Boston Children’s Hospital Intellectual and Developmental Disabilities Research Center Molecular Genetics Core Facility supported by U54HD090255 from the NIH Eunice Kennedy Shriver National Institute of Child Health and Human Development. We also thank the brain tissue donors and their families, from both the UMB Neurobiobank and Autism BrainNet. Autism BrainNet is a resource of the Simons Foundation Autism Research Initiative (SFARI). Autism BrainNet also manages the Autism Tissue Program (ATP) collection, previously funded by Autism Speaks. We are grateful and indebted to the families who donated tissue for research purposes to Autism BrainNet and the ATP. Supported by the NIMH (U01MH106883) and the Tan Yang Center for Autism Research at Harvard Medical School. C.A.W. is an Investigator of the Howard Hughes Medical Institute. S.K.A. is supported by a Paul and Daisy Soros Fellowship for New Americans. C.M.D. was supported in part by NIMH Translational Post-doctoral Training Program in Neurodevelopment T32MH112510.

## Contributions

C.M.D. performed the experiments, analyzed the data, and wrote the manuscript draft. M.T. and K.W. contributed to single-nuclei analysis. S.K.A. performed experiments. C.A.W conceptualized and supervised the project. All authors reviewed and contributed to revising and editing the manuscript draft.

---

**Figure 2 figure supplement 1:**
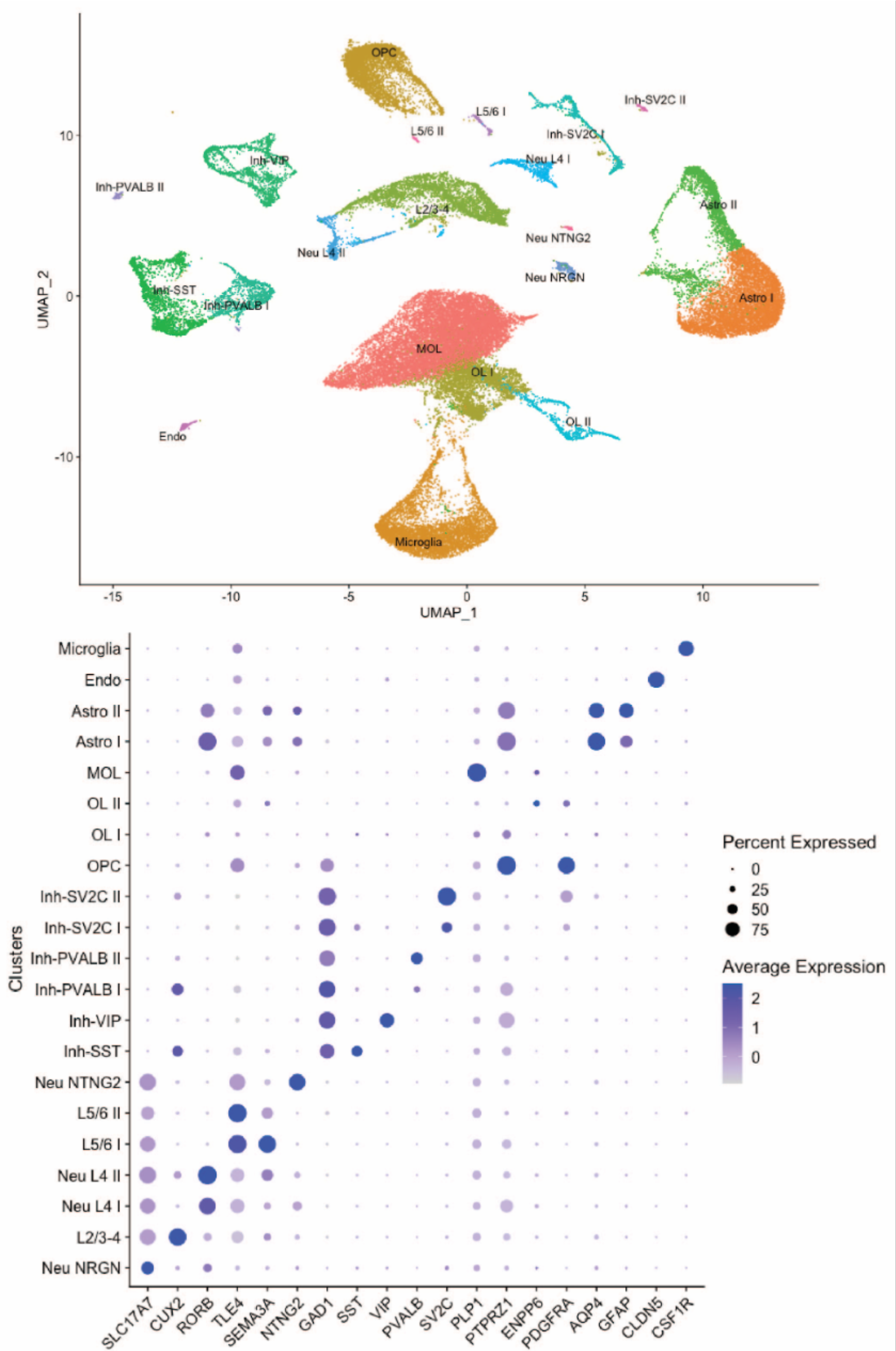
High resolution clustering of frontal cortex corroborates more general classification. Nuclei clustering of frontal cortex reveals accurate assignment of inhibitory neuronal subclusters and layer specific markers. Abbreviations as per Figure 2. Top panel is integrated UMAP, bottom is dot plot of cell type and layer specific marker expression by cluster.

**Figure 2 figure supplement 2:**
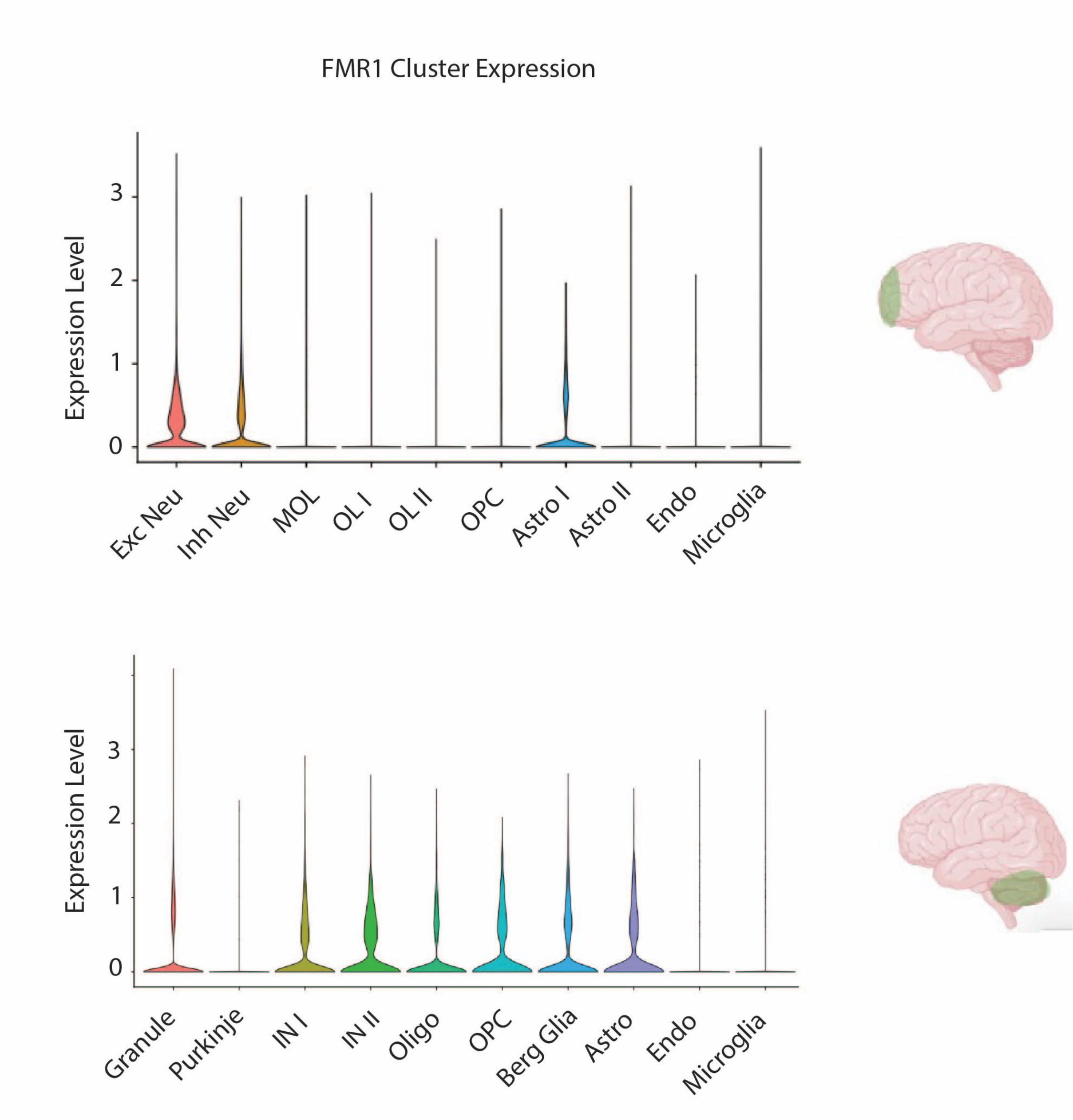
Average FMR1 expression by cluster and region. Abbreviations as per Figure 2.

**Figure 2 figure supplement 3:**
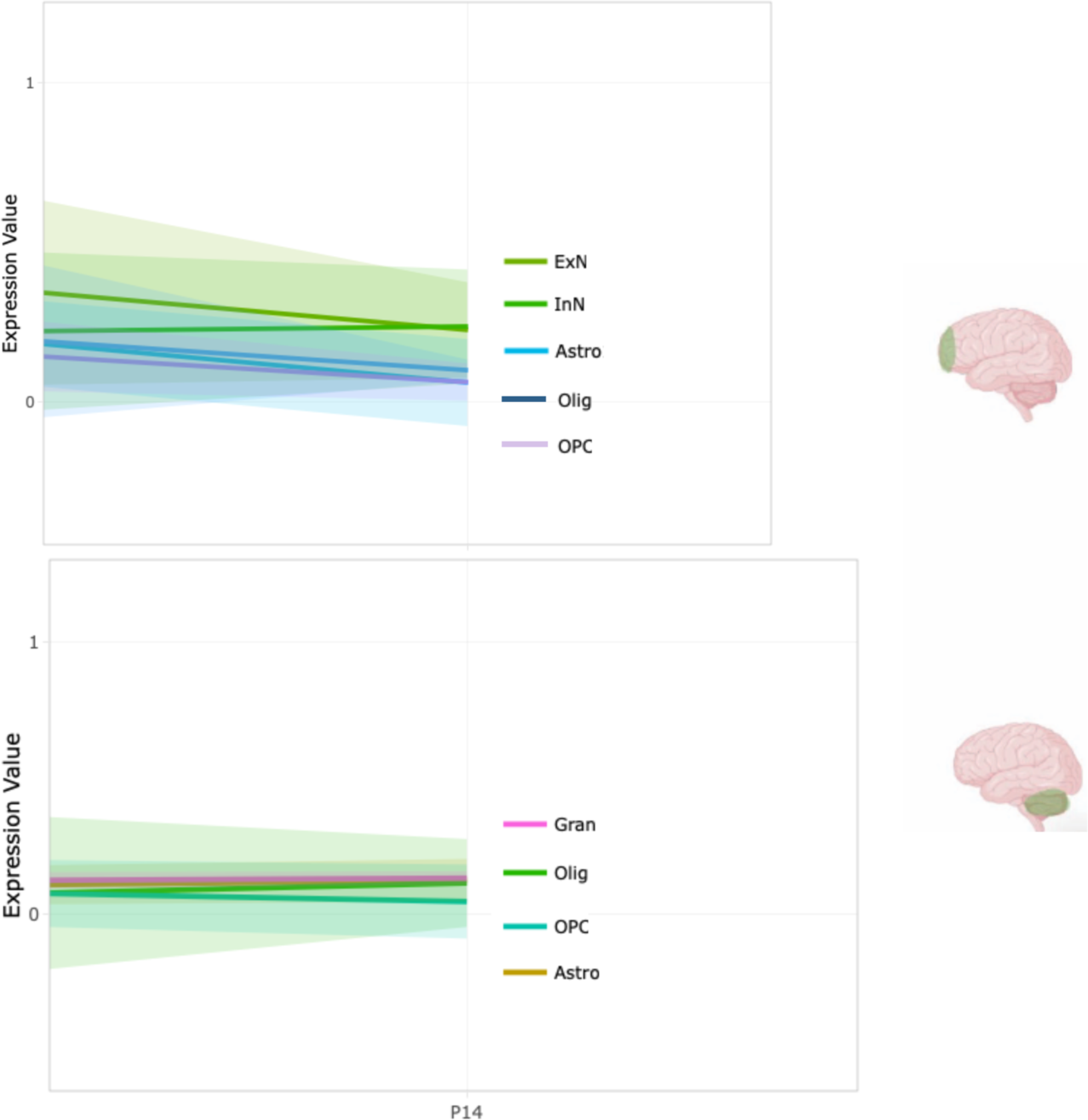
FMR1 expression in human frontal cortex (top) and cerebellum (bottom) in independent scRNA dataset replicates higher cerebellar glial expression P14: ages 40-59. Generated from stab.comp-sysbio.org on 1/8/21. Additional cell types hidden for clarity of presentation.

**Figure 2 figure supplement 4:**
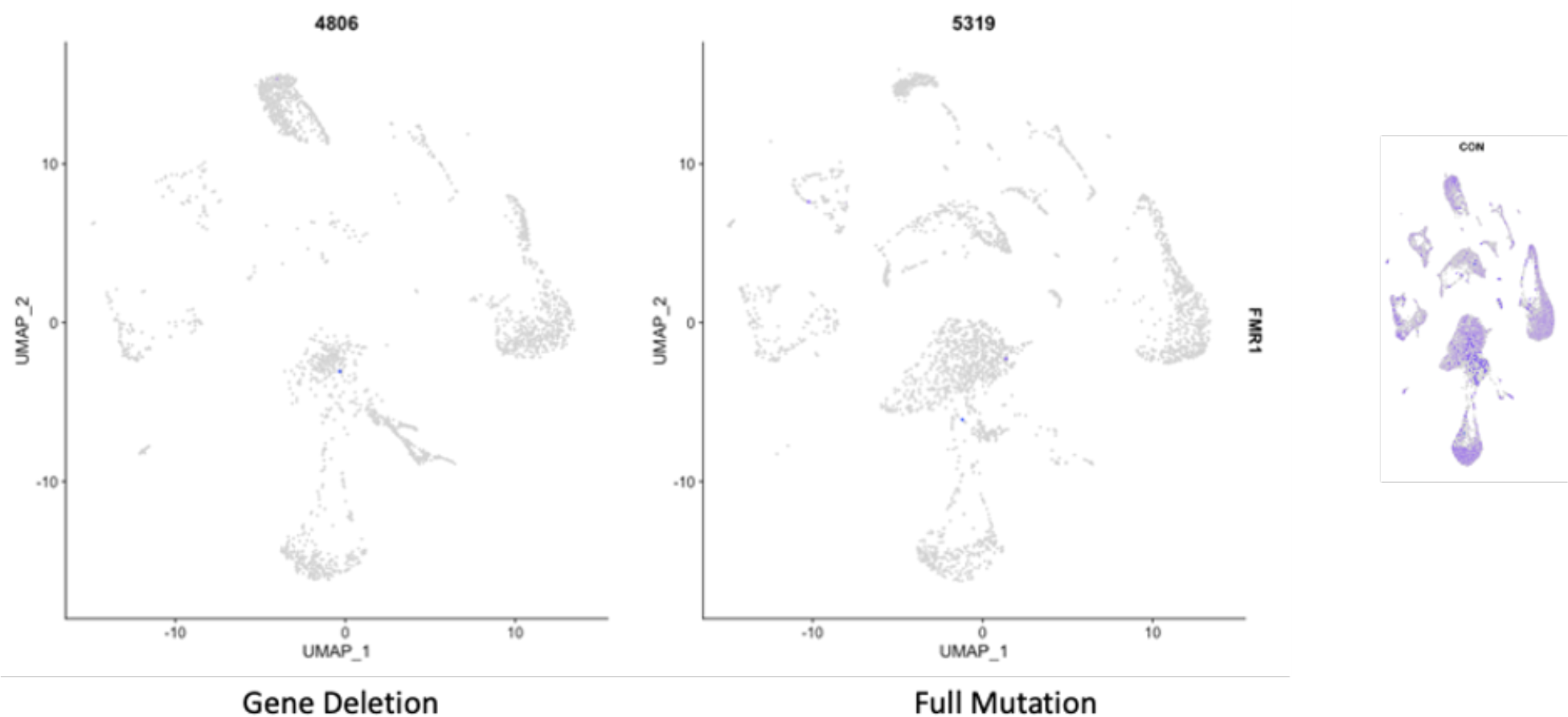
*FMR1* expression is eliminated in all cell clusters in Fragile X syndrome, regardless of genetic mechanism. Expected expression (purple) presented in right panel demonstrates average control expression. Sort cell feature in Seurat applied to visualize loss of expression. Frontal cortex UMAP is presented, results were identical in cerebellum.

**Figure 3 figure supplement 1:**
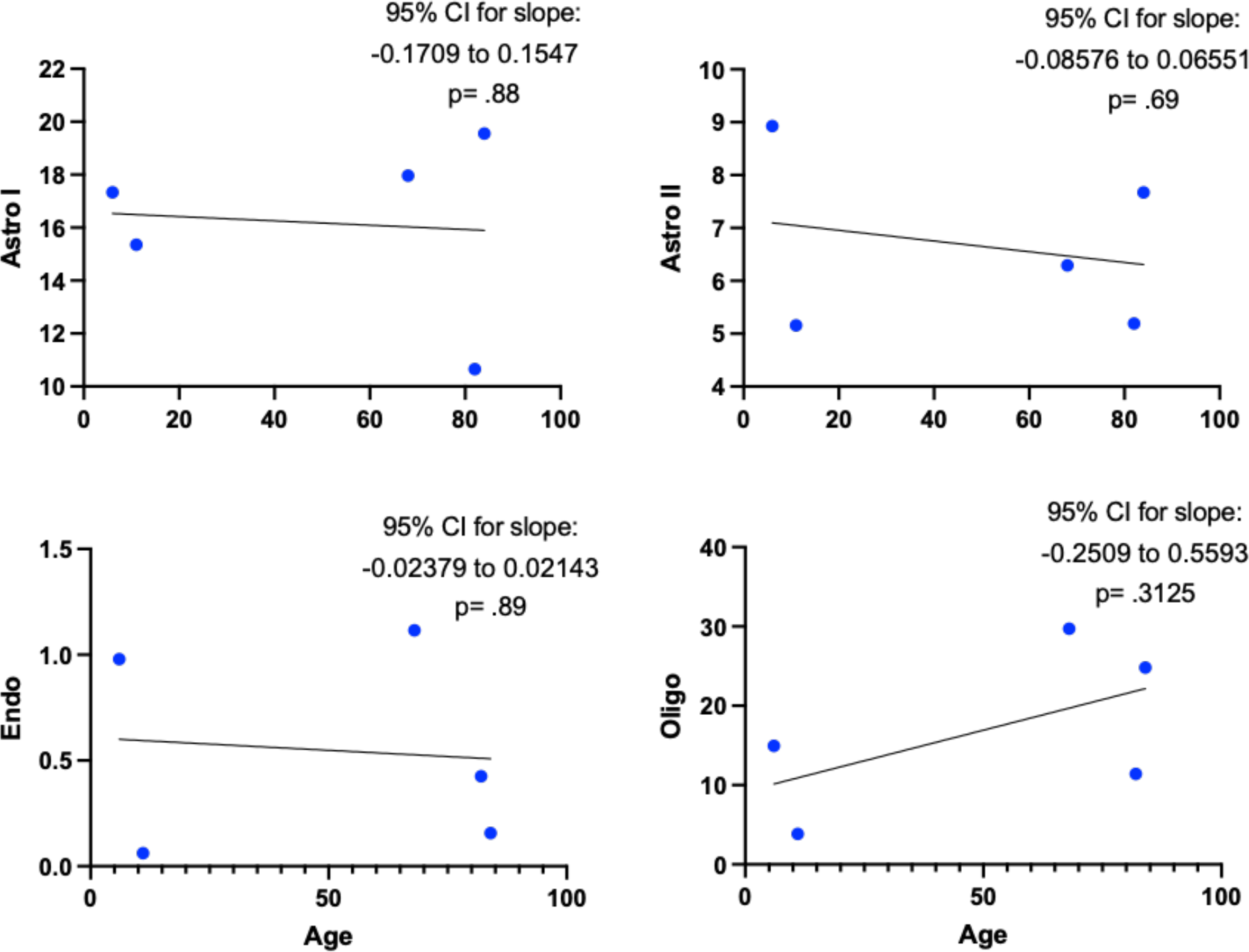
Linear regression for cluster proportion and age revealed no significant association with age for glial subclusters. Pvalue on graph for non-zero slope. Astro- astrocyte, Endo- endothelial, Oligo-mature oligodendrocyte (MOL).

**Figure 3 figure supplement 2:**
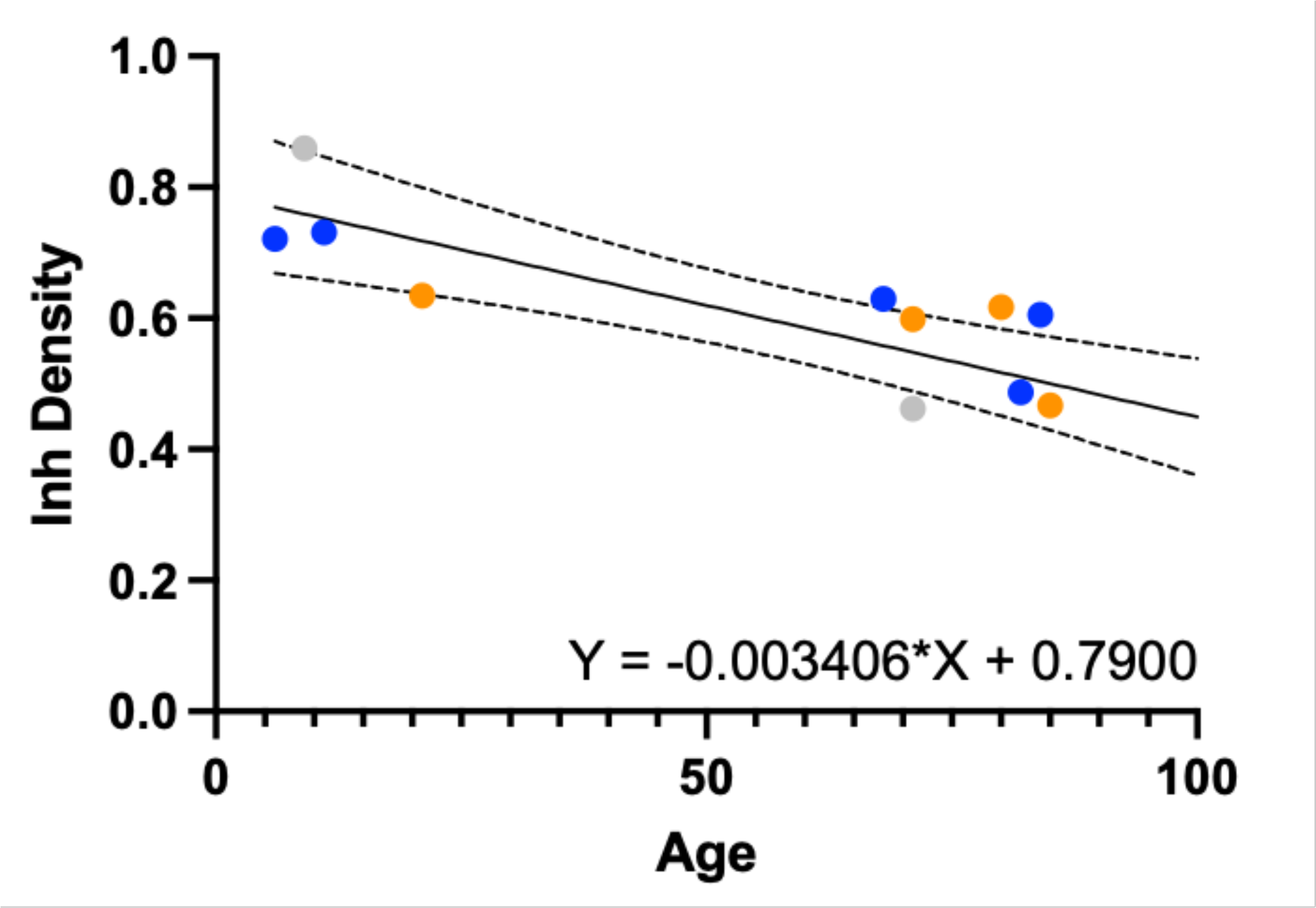
Linear regression for inhibitory/total neuron density in frontal cortex reveals significant age related decline that is not related to Fragile X status. Grey- FXS, blue- control, orange- PM. p-value of non-zero slope < .001.

**Figure 3 figure supplement 3:**
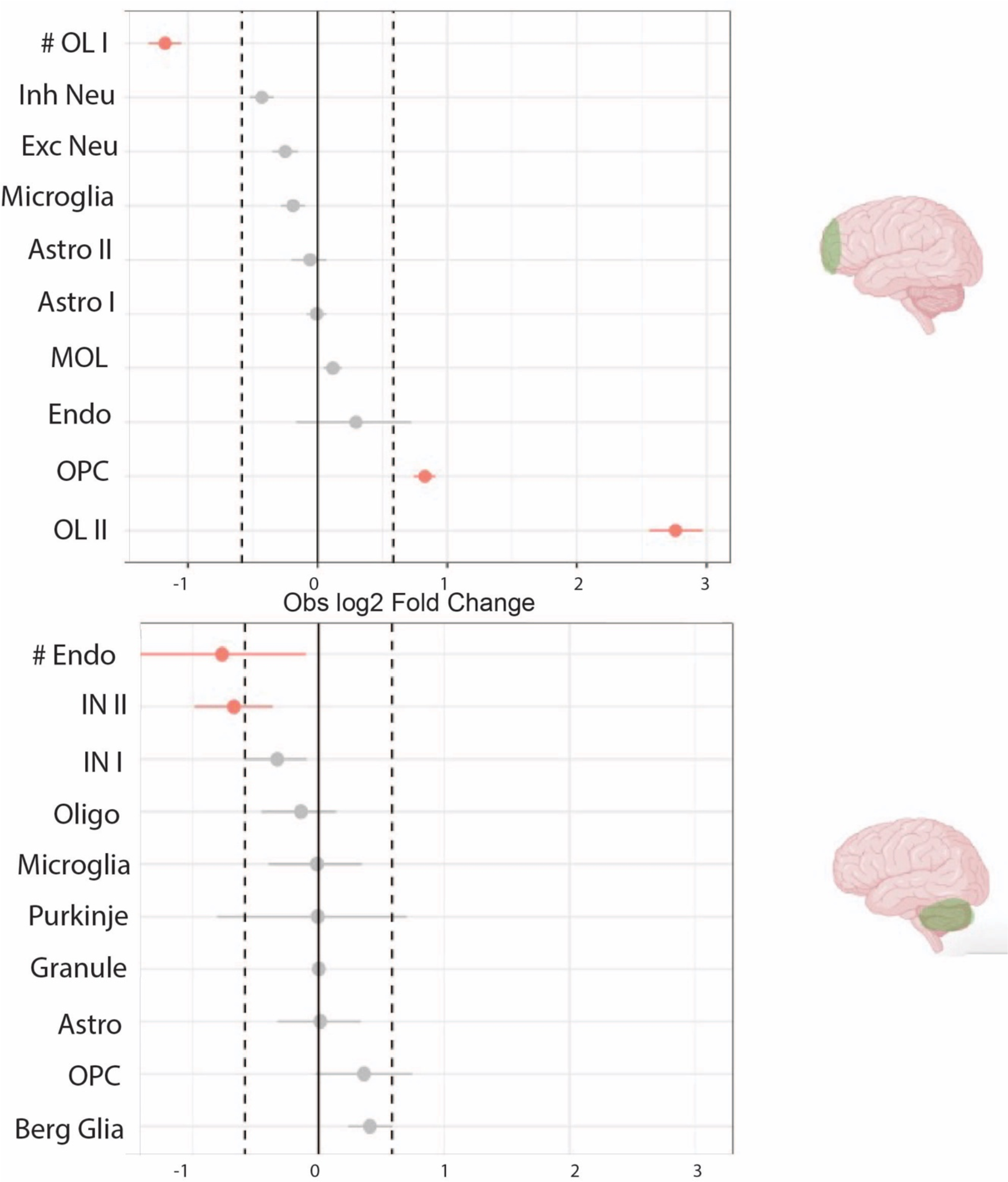
Alterations in cellular composition in Fragile X syndrome. Permutation plot resulting from scProportion() with red indicating FDR < .05 and abs(log2FC) > 0.58. # indicates significance of this cluster is not robust to outlier sample removal.

**Figure 4 figures supplement 1:**
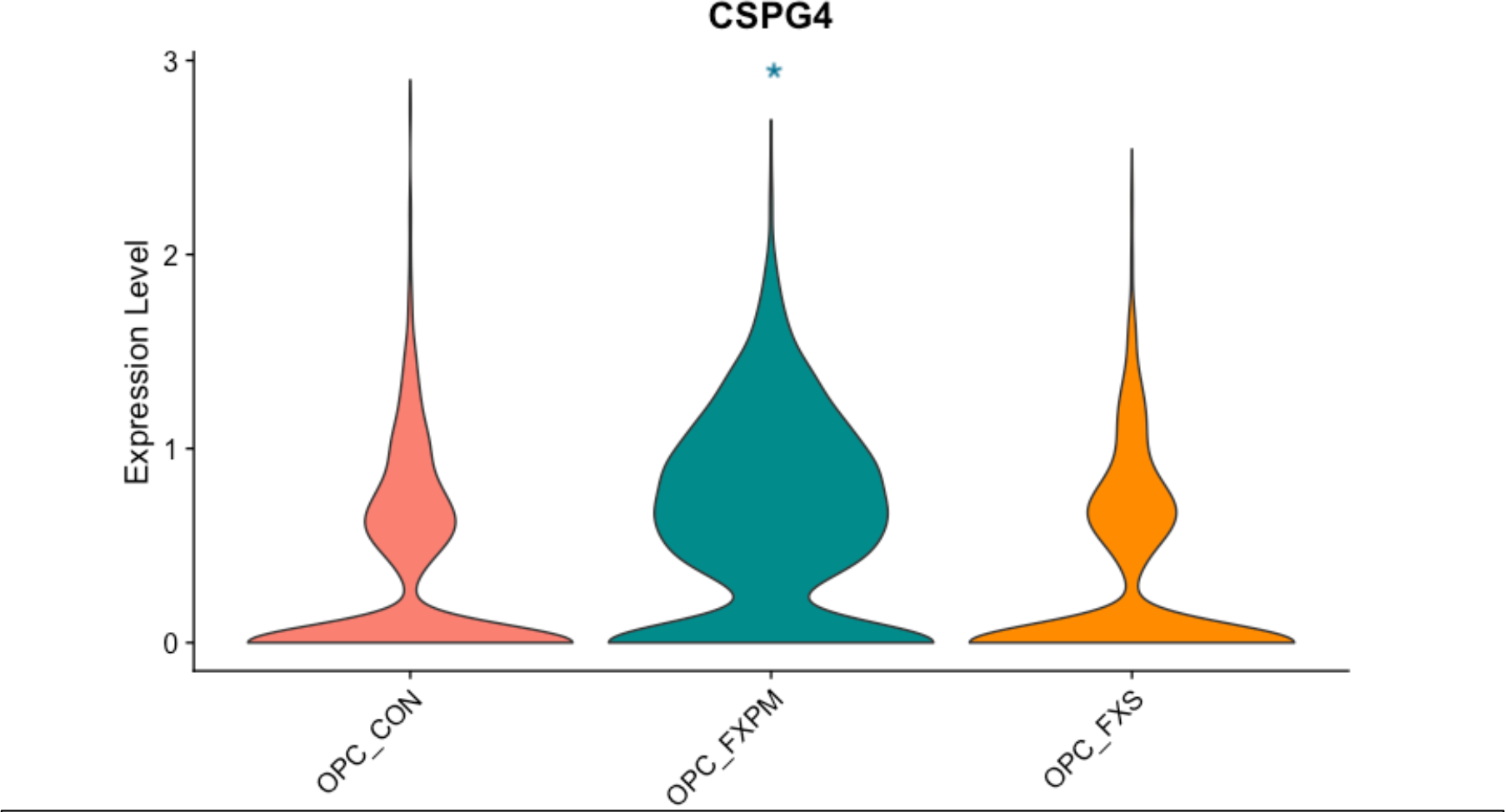
Increased CSPG4 (NG2) expression in OPCs in PM cases in frontal cortex. * indicates comparison vs control, padj <.05.

**Figure 4 figure supplement 2:**
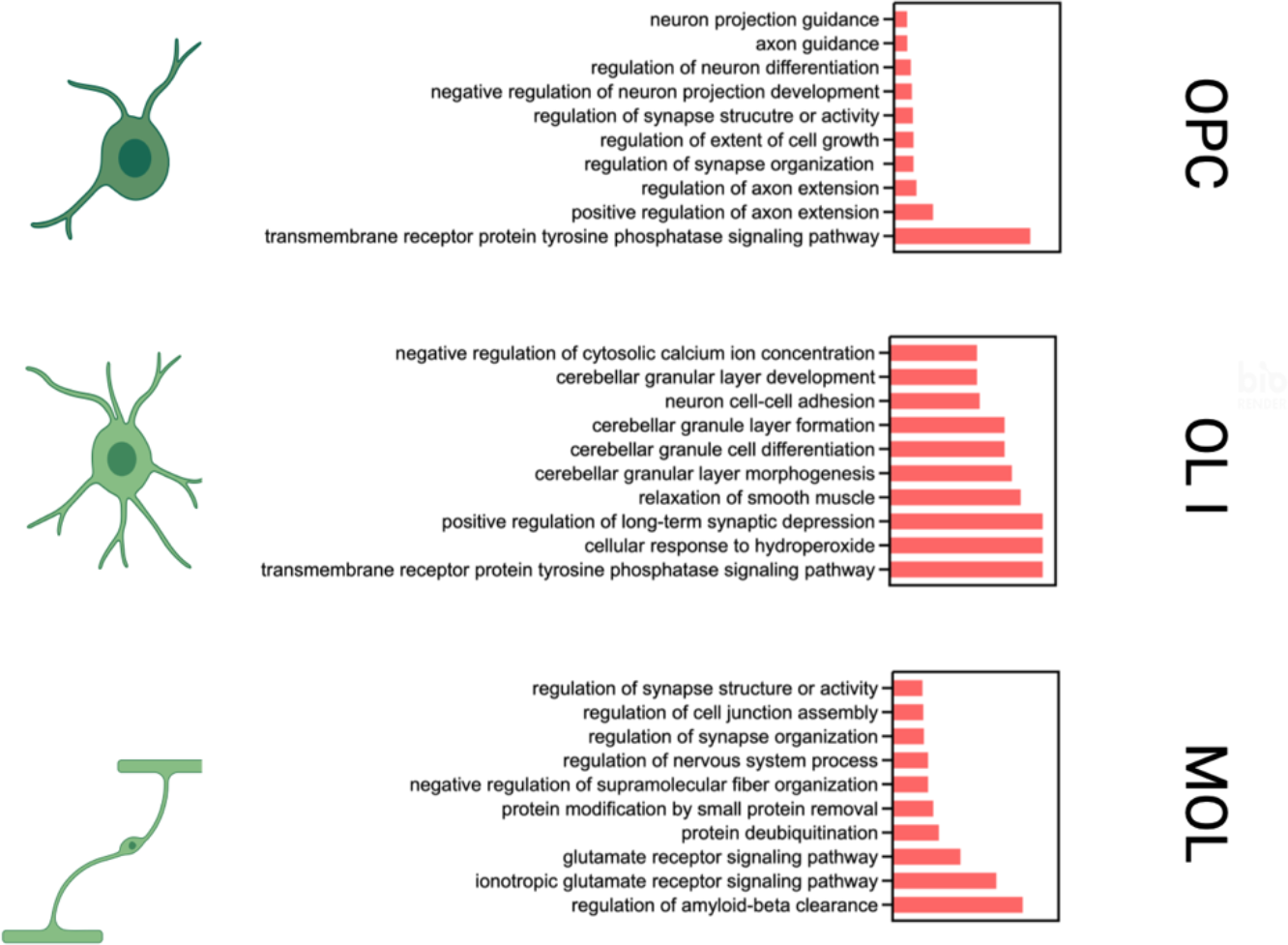
GO analysis of downregulated genes in oligodendrocyte clusters in PM cases. Analysis reveals disturbances in synaptic and neuronal organization and regulation.

**Figure 4 figure supplement 3:**
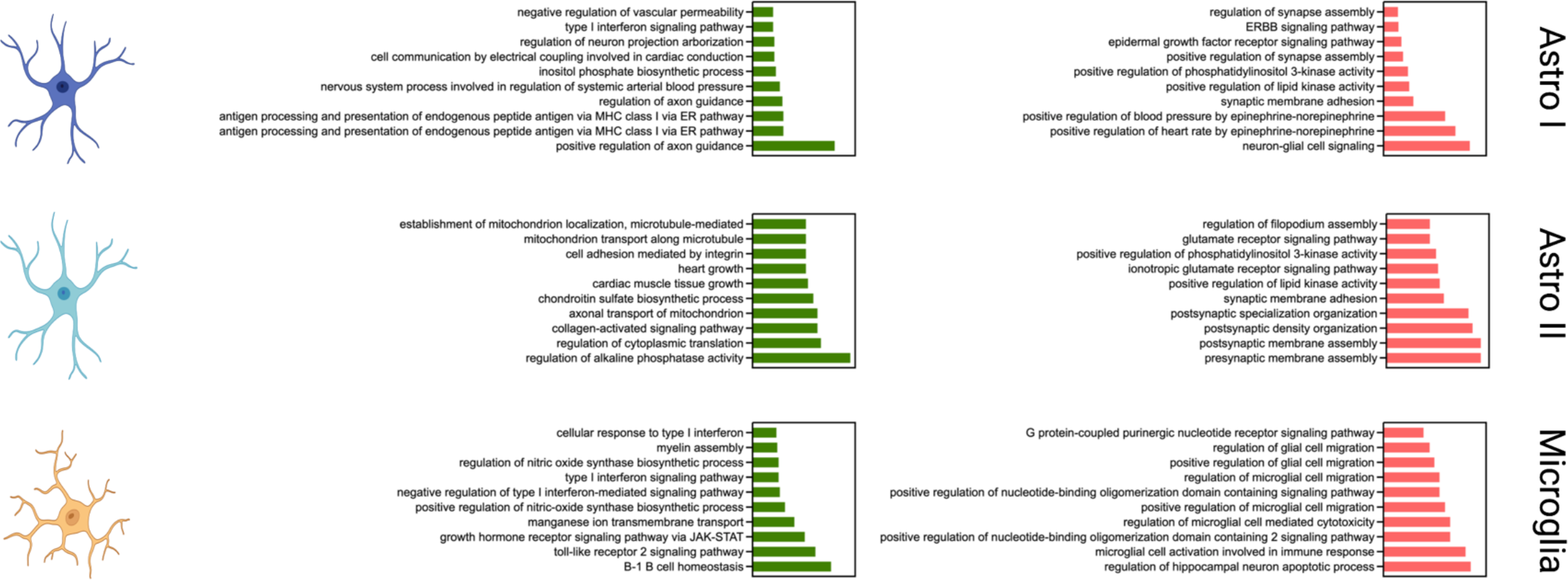
GO analysis of upregulated (green) and downregulated (red) genes in astrocyte and microglia clusters in PM cases. Analysis reveal upregulation of inflammation as well as disturbances in synaptic and neuronal organization and regulation.

**Figure 4 figure supplement 4:**
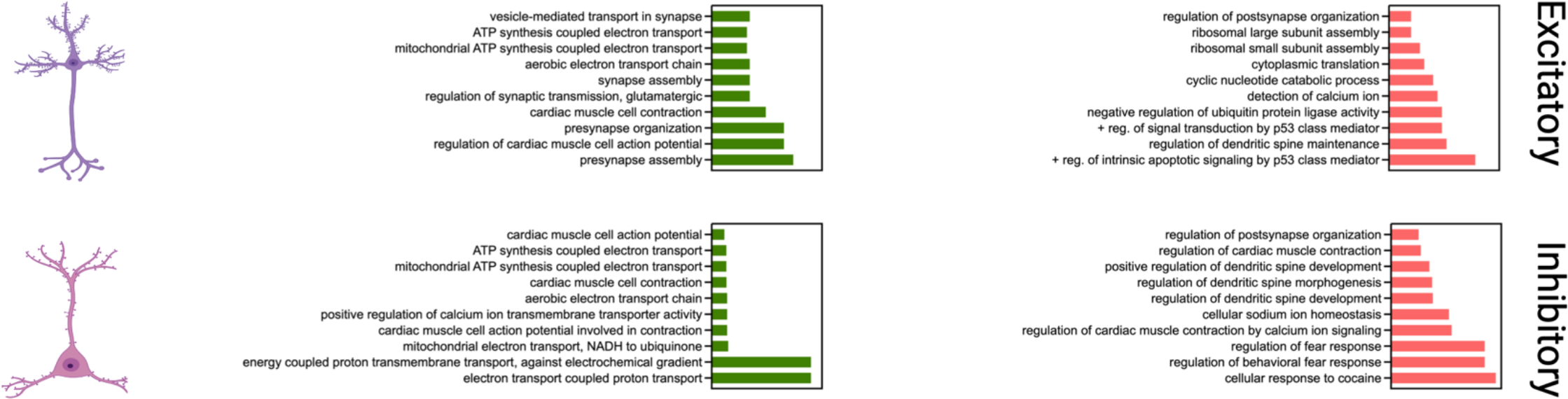
GO analysis of upregulated (green) and downregulated (red) genes in neuronal clusters in PM cases. There is evidence of metabolic stress and downregulation of synaptic and dendritic organization and development.

**Figure 4 figure supplement 5:**
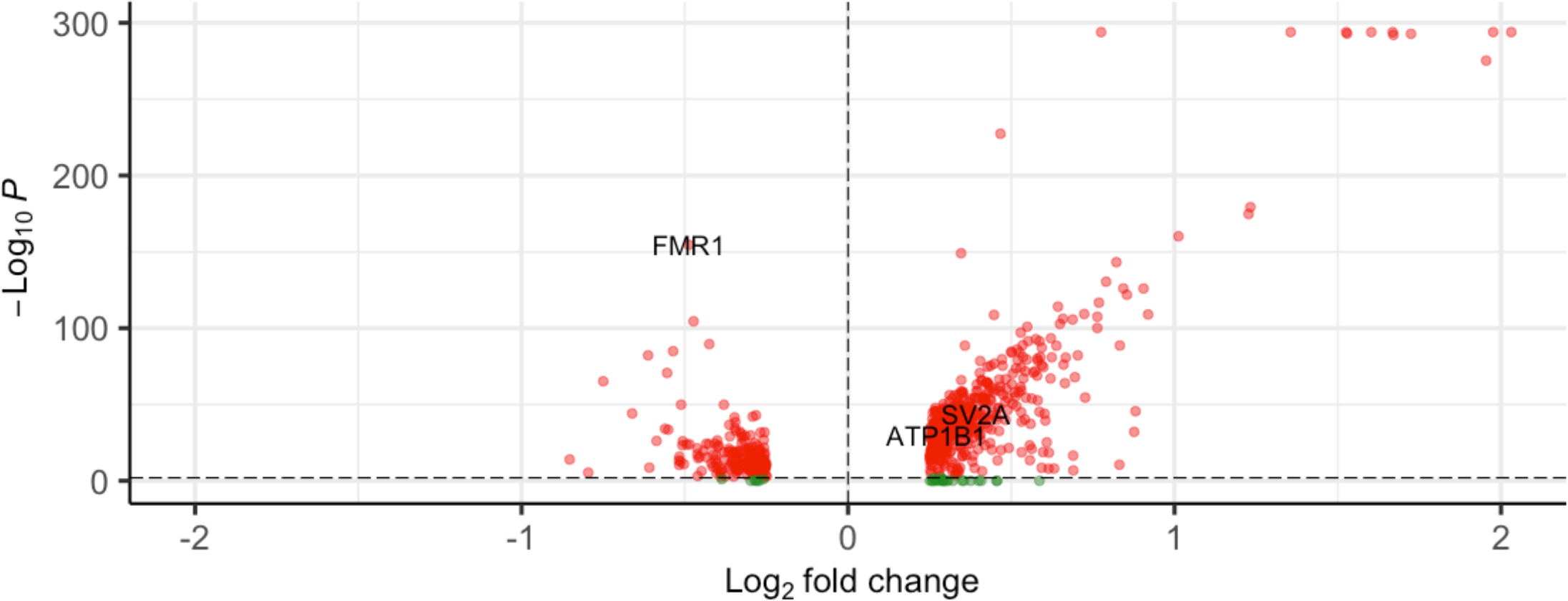
Volcano plot of differentially expressed genes in FXS in frontal cortex excitatory neurons. Note expected loss of FMR1 and deregulation of select FMRP target genes (ATP1B1, SV2A).

**Figure 4 figure supplement 6:**
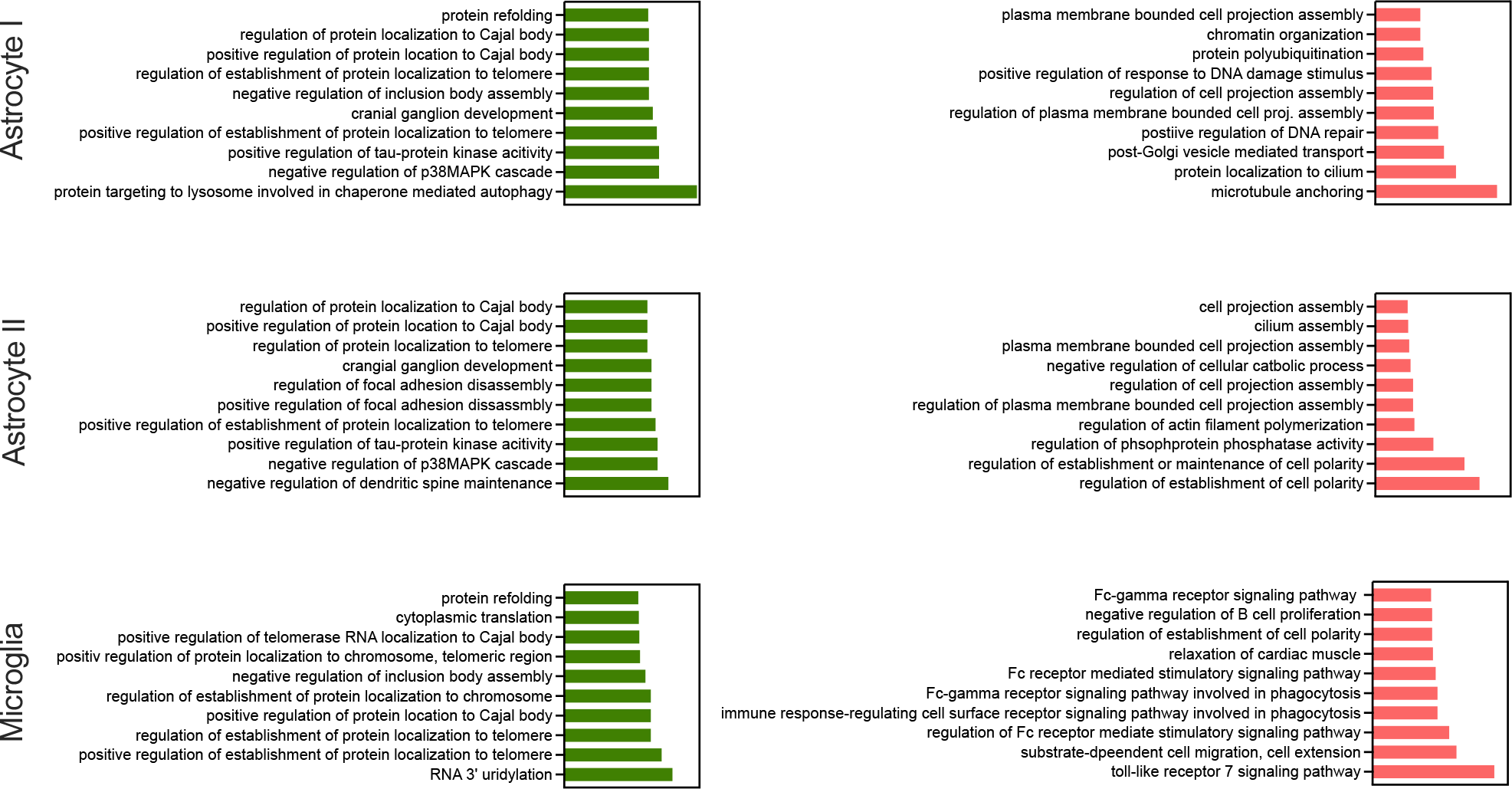
GO analysis of upregulated (green) and downregulated (red) differentially regulated genes in astrocytes and microglia in FXS.

**Figure 4 figure supplement 7:**
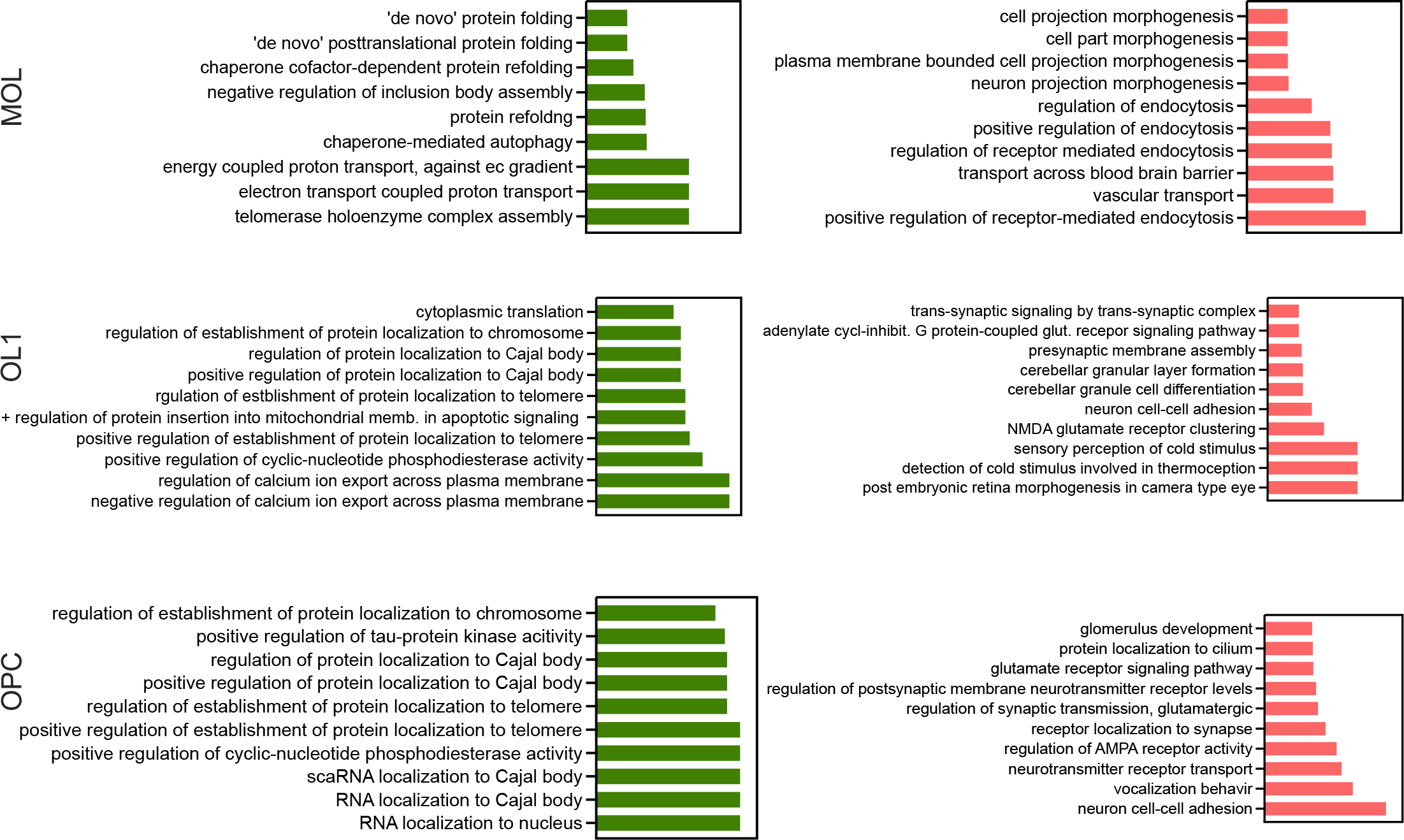
GO analysis of upregulated (green) and downregulated (red) differentially regulated genes in oligodendrocyte lineage in FXS.

**Figure 4 figure supplement 8:**
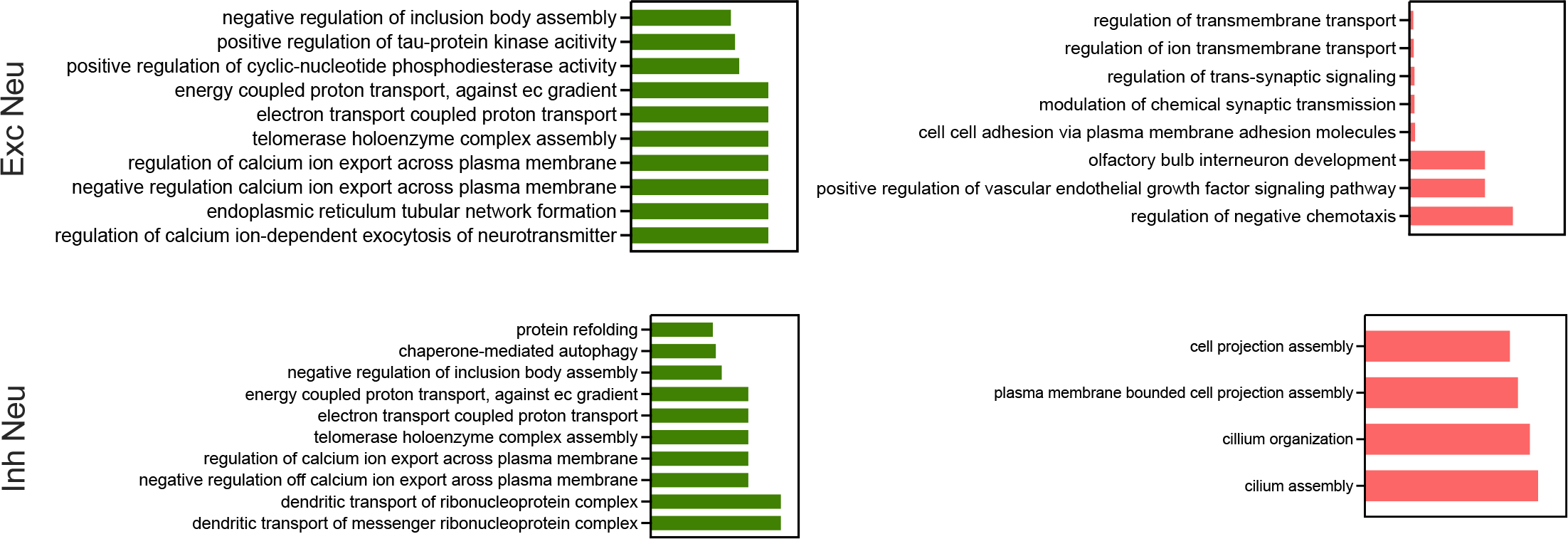
GO analysis of upregulated (green) and downregulated (red) differentially regulated genes in neurons in FXS.

**Figure 4 figure supplement 9:**
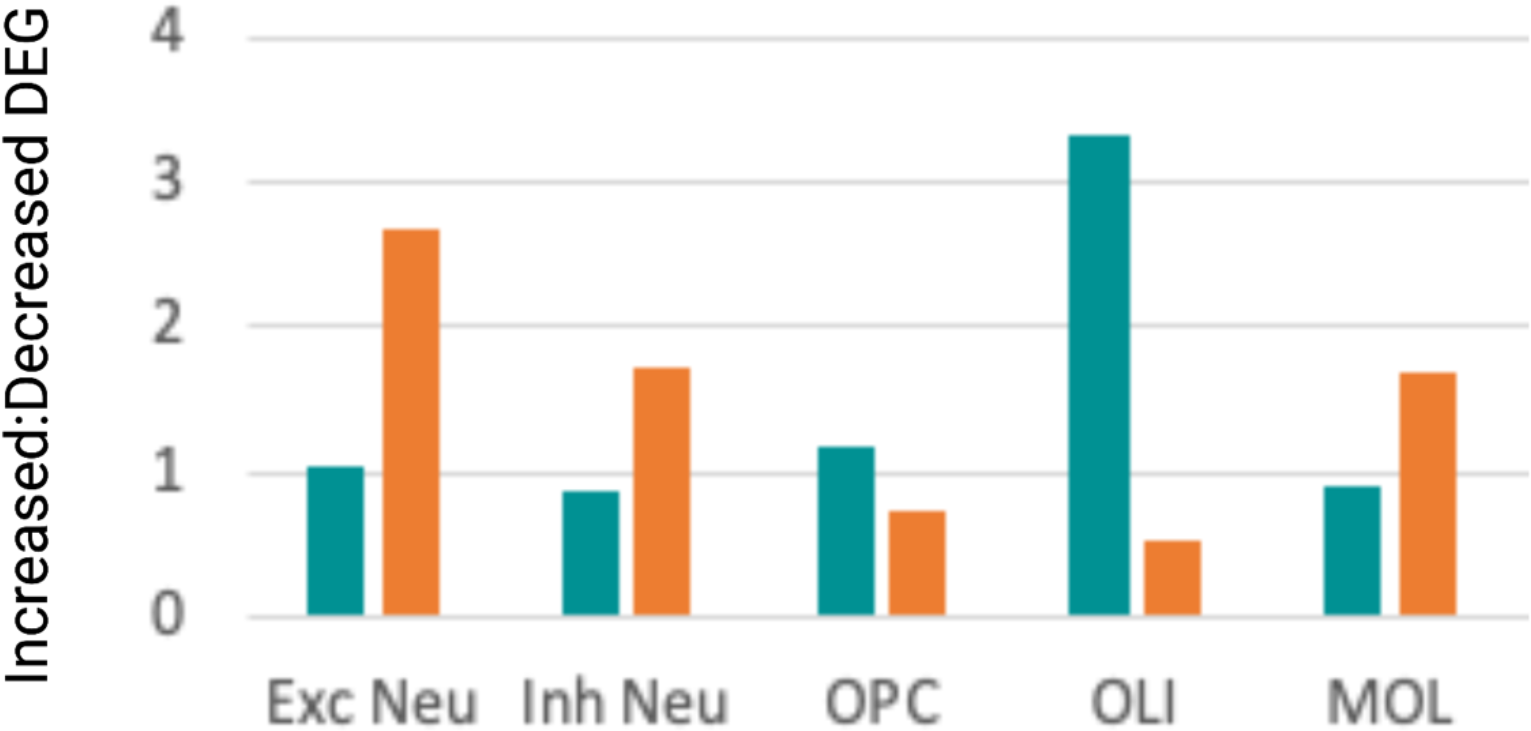
Comparison of ratio of increased:decreased differentially expressed genes in select clusters in PM cases (teal) and FXS (orange) reveal distinct patterns of global transcriptional dysregulation.

**Appendix Fig 1:**
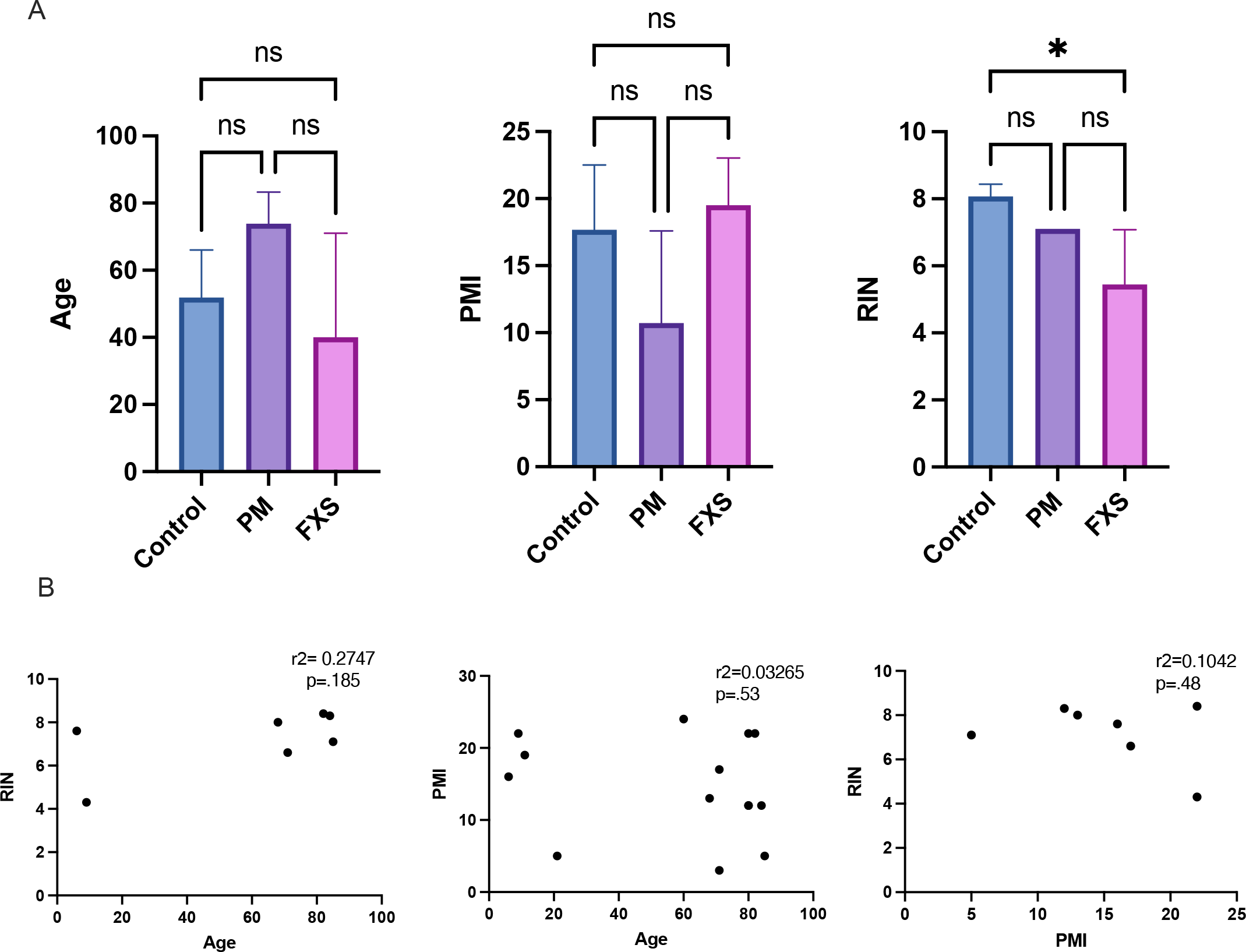
A. Age, PMI, and RIN by condition, ns= non-significant, one-way ANOVA, Tukey’s multiple comparison test, * p < .05. B. No significant association between RIN and age, PMI and age, and RIN and PMI.

**Appendix Fig 2:**
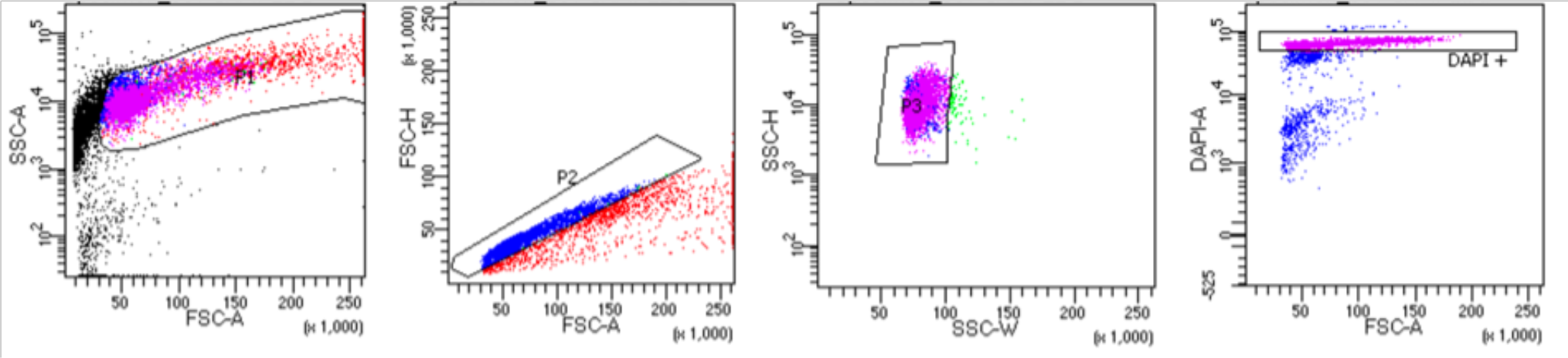
Representative flow cytometry results for nuclear selection- ∼ 5-10% parent population is selected.

